# Incorporation of multiple β^2^-backbones into a protein *in vivo* using an orthogonal aminoacyl-tRNA synthetase

**DOI:** 10.1101/2023.11.07.565973

**Authors:** Noah X. Hamlish, Ara M. Abramyan, Alanna Schepartz

## Abstract

Synthesis of sequence-defined biomaterials whose monomer backbones diverge from canonical α-amino acids represents the next frontier in protein and biomaterial evolution with the potential to yield better biological therapeutics, bioremediation tools, and biodegradable plastic-like materials. One monomer family of particular interest for biomaterials are β-hydroxy acids. Many natural products contain isolated β-esters, and polymeric β-esters are found in polyhydroxyalkanoate (PHA) polyesters under development as bioplastics and drug encapsulation/delivery systems. Here we report that β^2^-hydroxy acids possessing both *(R)* and *(S)* absolute configuration are excellent substrates for pyrrolysyl-tRNA synthetase (PylRS) enzymes *in vitro*, and that *(S)*-β^2^-hydroxy acids are substrates *in cellulo*. Using the *Ma*PylRS/*Ma*tRNA^Pyl^ pair, in conjunction with wild-type *E. coli* ribosomes and EF-Tu, we report the cellular synthesis of model proteins containing two *(S)*-β^2^-hydroxy acid residues at internal positions. Metadynamics simulations provide a rationale for the observed enantioselective preference of the ribosome for the *(S)*-β^2^-hydroxy acid backbone and mechanistic insights that inform future ribosomal engineering efforts. As far as we know, this finding represents the first example of an orthogonal synthetase that accepts a β-backbone substrate and the first example of a protein hetero-oligomer containing multiple expanded-backbone monomers produced *in cellulo*.

## Introduction

There is great interest in the synthesis and study of sequence-defined biomaterials whose monomer backbones diverge from canonical α-amino acids. Such hetero-oligomers can adopt new and known secondary and tertiary structures^1–5^ that confer novel functions including enhanced proteolytic resistance^6–8^ and membrane permeability.^9,10^ As a class, sequence-defined biomaterials provide otherwise non-existent opportunities to expand and evolve protein and biomaterial structure and function and develop improved biological therapeutics.^11,12^

One monomer family of particular interest for biomaterials consists of β-hydroxy acids (β-HAs) (**Figure 1A**). β-HAs embody both an expanded backbone and a non-proteinogenic nucleophile and assemble into biomaterials known as β-esters. Isolated β-esters are found in therapeutically relevant natural products (enterobactin, previoprolide),^13^ biosurfactants with environmental applications (surfactin),^14^ and FDA-approved therapeutics (romidepsin, rapamycin) (**Figure 1B**).^15,16^ Polymeric β-esters are naturally found in polyhydroxyalkanoate (PHA) polyesters, which are currently in development as bioplastics^17^ and drug encapsulation/delivery systems.^18^

**Figure 1.**
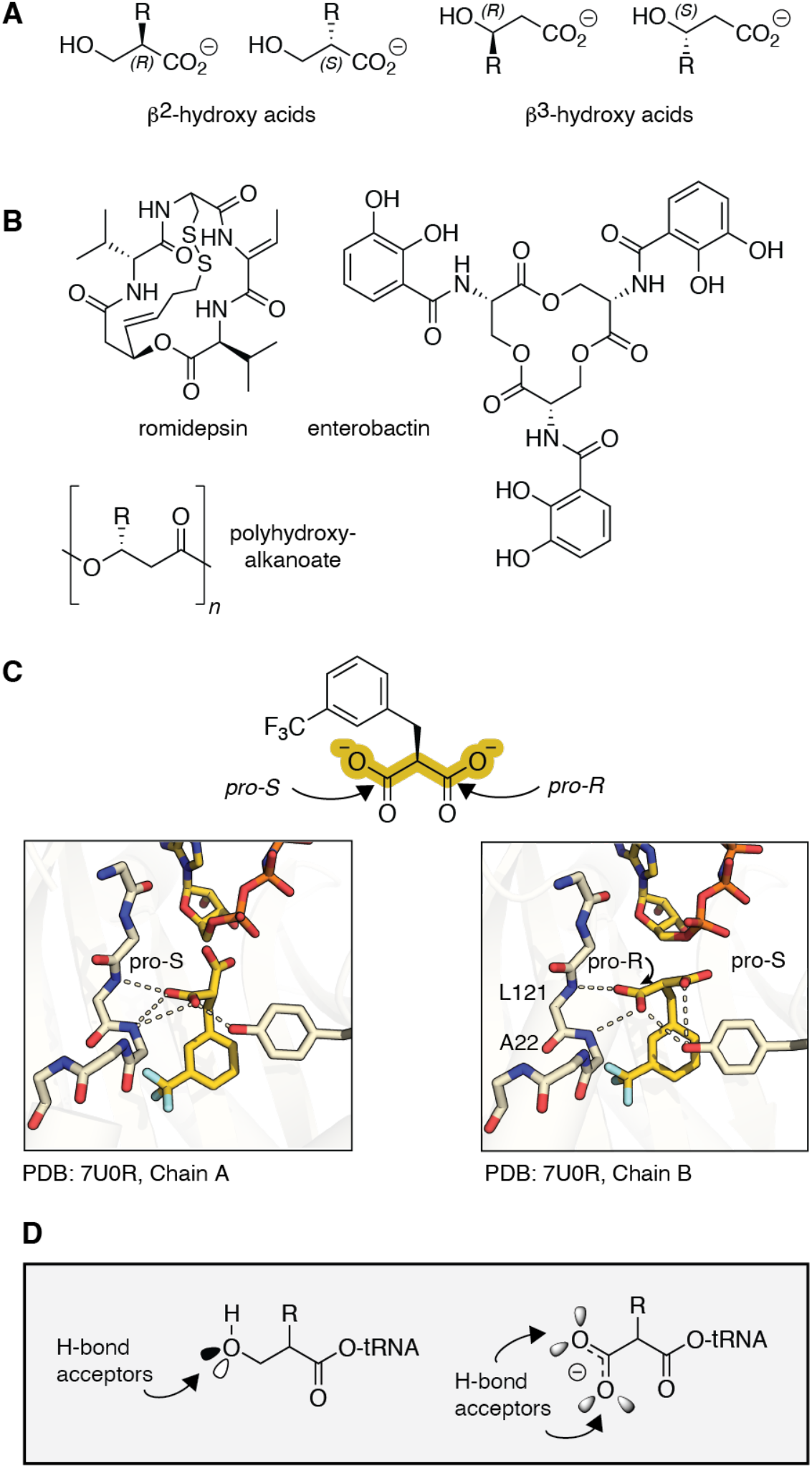
β-hydroxy acids in natural products and as substrates for pyrrolysyl-tRNA synthetase (PylRS) variants. (A) Structures of β^2^- and β^3^-hydroxy acids and (B) natural products and biomaterials that contain β^2^- or β^3^-hydroxy esters. (C) Structure of *Ma*FRSA bound to *m*-CF_3_-2-benzylmalonate (PDB 7U0R) illustrates the absence of a bound active site water and direct H-bonds from L121 and A122 (*Ma* numbering). (D) This work tests the hypothesis that β^2^-hydroxy acids will also act as substrates for PylRS enzymes by virtue of their ability to accept one or two backbone H-bonds.

Although there exists one report in which a single β^2^-ester has been introduced into a protein using the wild-type *E. coli* ribosome *in vitro*,^19^ there are no examples in which any β^2^-ester has been introduced into a ribosomal product *in vivo*. Indeed, there remains only one example^20^ in which a β-backbone of any type has been introduced into a protein in a cell, and that effort required the endogenous and non-orthogonal *E. coli* PheRS synthetase and an engineered ribosome with a structurally characterized assembly defect.^20,21^ One challenge limiting the *in vivo* ribosomal synthesis of sequence-defined β-ester biomaterials is the absence of an orthogonal aminoacyl-tRNA synthetase (aaRS)/tRNA pair that accepts a β-HA as a substrate.

Previous work has shown that the widely employed and orthogonal pyrrolysyl-tRNA synthetase (PylRS)^22^, as well as an engineered derivative *Ma*FRSA,^23^ accept several non-α-amino acid substrates, including those in which the α-NH_2_ group is replaced with α-H, α-OH, α-SH, α-*N*-methyl, α-*N*-formyl, and α-carboxy substituents.^24–27^

Examination of the structure of *Ma*FRSA bound to one such α-carboxy substrate, *m*-CF_3_-2-benzylmalonate, provided two insights into how PylRS enzymes might engage expanded backbone monomers such as a β^2^-HA. First, although the structure of PylRS bound to pyrrolysine shows the substrate α-amine coordinated to the homodimeric enzyme in a single conformation *via* a well-defined active site water molecule,^28^ that water is absent in the structure of *Ma*FRSA bound to *m*-CF_3_-2-benzylmalonate (**Figure 1C**), presumably to accommodate the expanded size of the α-substituent. In the absence of this bound water, new amide backbone H-bonds from residues L121 and A122 engage the α-carboxy group instead of the bound water. Second, in the structure of FRSA bound to *m*-CF_3_-2-benzylmalonate, the two subunits of the dimeric enzyme bind the prochiral *m*-CF_3_-2-benzylmalonate substrate in stereochemically distinct configurations. In one active site, the *pro-S* carboxylate of *m*-CF_3_-2-benzylmalonate engages the backbone amides of L121 and A122; in the other, the substrate rotates and the *pro-R* carboxylate is coordinated instead (**Figure 1C**). These results imply that the backbone H-bonds from residues L121 and A122 are highly stabilizing, and PylRS can accommodate substrates retaining the same absolute configuration as natural L-α-amino acids as well as their enantiomers.

Like the α-carboxy group of a malonic acid, a β^2^-hydroxyl group of tRNA acylated with a β^2^-HA can also accept at least one, and perhaps two H-bonds from the synthetase amide backbone (**Figure 1D**). Here we report that β^2^-hydroxy acids possessing both *(R)* and *(S)* absolute configurations are substrates for PylRS enzymes *in vitro*. Further, we report the unexpected finding that only *(S)*-β^2^-hydroxy acids–whose absolute configuration maps onto a D-α-amino acid–are substrates *in cellulo*. Using the *Ma*PylRS/*Ma*tRNA^Pyl^ pair in classic expression strains (BL21 and C321.ΔA.exp), we report the cellular synthesis of model proteins containing two *(S)*-β^2^-HA residues at internal positions. Metadynamics simulations provide a clear rationale for the observed enantioselective preference for the *(S)*-β^2^-hydroxy acid backbone and provide mechanistic insights useful for future ribosomal engineering efforts. As far as we know, this finding represents the first example of an orthogonal synthetase that accepts a β-backbone substrate and the first example of a protein hetero-oligomer containing multiple expanded backbone monomers produced *in cellulo*.

## Results

### *Ma*PylRS accept β^2^-hydroxy acids as substrates *in vitro*

To assess if PylRS-like enzymes would accept β^2^-hydroxy acids as substrates, we performed *in vitro* tRNA acylation reactions using purified enzymes and directly analyzed the products using intact tRNA LC-HRMS (Figure 2A).^25,29^ We began with *Ma*PylRS and compared the yields of acylated tRNA^Pyl^ from reactions containing the known substrates *(S)*-α-NH_2_-N^ε^-Boc-Lysine (*(S)*-α-ΝΗ_2_ **1**) and *(S)*-α-OH-N^ε^-Boc-Lysine (*(S)*-α-OH **2**) as well as the enantiopure β^2^-OH analogs **3** and **4**. Reactions performed using 12.5 μM *Ma*PylRS, 25 μΜ *Ma*tRNA^Pyl^, and 10 mM **1** or **2** and incubated for 2 hours at 37°C generated the expected mono-acylated tRNA products **1**-acyl-tRNA^Pyl^ (23188.4 Da) and **2**-acyl-tRNA^Pyl^ (23189.6 Da) (Figure 2B**, Supplementary** Figure 1) in yields of 56% and 59%, respectively (Figure 2C). As observed previously,^25^ *Ma*PylRS-promoted acylation of tRNA^Pyl^ with *(S)*-α-OH **2** also generated a di-acylated tRNA^Pyl^ product (Figure 2C). Analogous reactions supplemented with *(S)*-β^2^-OH **3** or *(R)*-β^2^-OH **4** also generated the expected diastereomeric mono-acylated tRNA products **3**-acyl-tRNA^Pyl^ and **4**-acyl-tRNA^Pyl^ (23203.6 Da) in yields of 27% and 79%, respectively (Figure 2B). Again, under these conditions both substrates also generated detectable levels of di-acylated tRNA^Pyl^; 68% for *(S)*-β^2^-OH **3** and 11% for *(R)*-β^2^-OH **4** (Figure 2C). Thus, under these conditions, both β^2^-OH acids **3** and **4** are substrates for *Ma*PylRS, as anticipated by the FRSA-bound structure of the pro-chiral substrate m-CF_3_-2-benzylmalonate (Figure 1C and D).^25^

**Figure 2.**
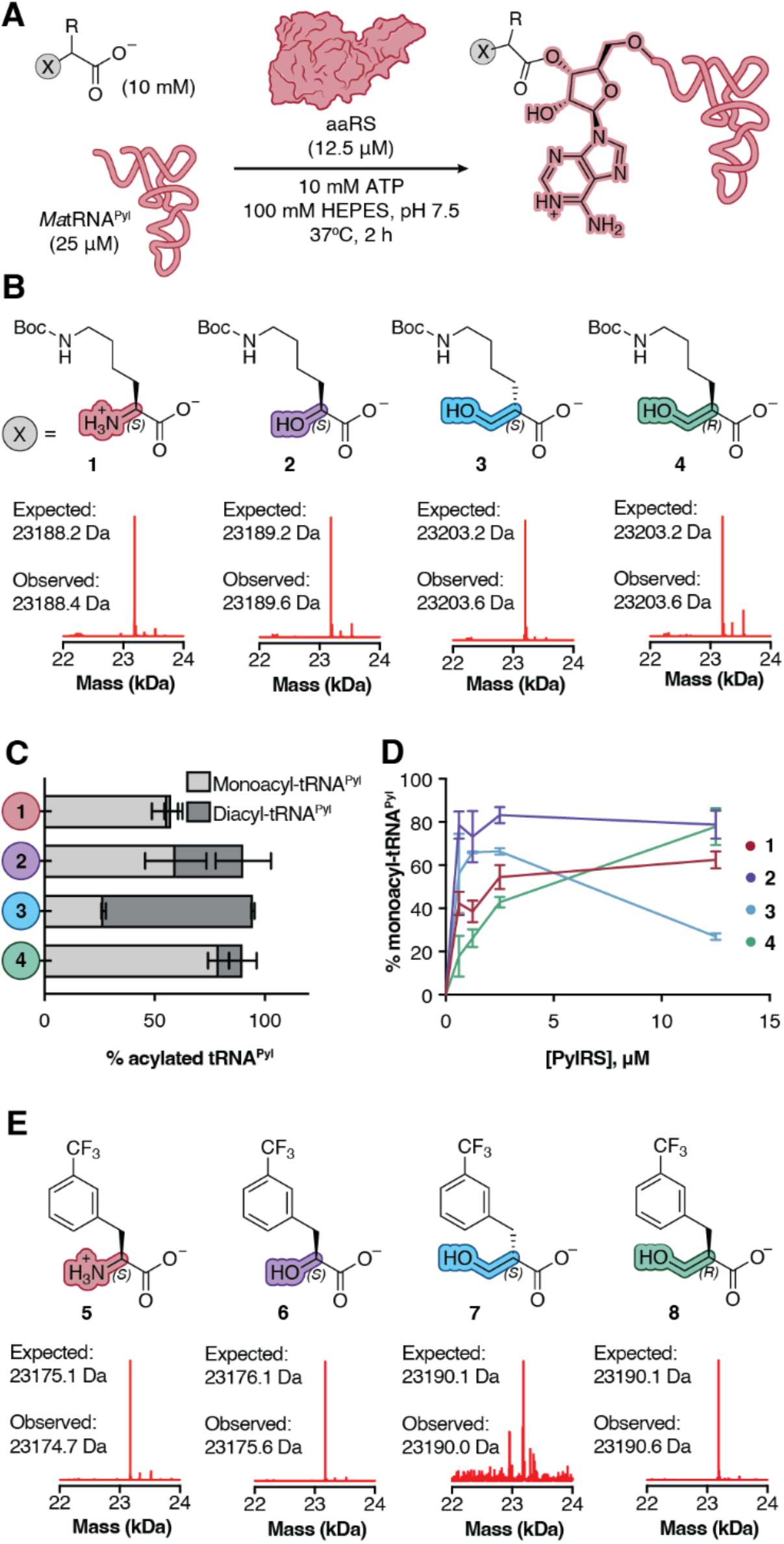
*Ma*PylRS and *Ma*FRSA accept β^2^-hydroxy acids as substrates *in vitro*. (A) Workflow for and (B) deconvoluted mass spectra of *in vitro* tRNA^Pyl^ acylation reactions containing 12.5 µM *Ma*FRSA and supplemented with monomers **1-4**. Signal is normalized to the highest signal in respective traces. Expected masses shown correspond to mono-acylated products. (C) Plot illustrating the relative yields of mono- and diacylated tRNA^Pyl^ generated during *in vitro* acylation reactions containing 12.5 μM PylRS and supplemented with 10 mM of monomer **1**, **2**, **3**, or **4**. (D) Plot illustrating yield of mono-acylated tRNA^Pyl^ as a function of [*Ma*PylRS].

It has been estimated that the concentration of a single aaRS enzyme expressed from an endogenous promoter in *E. coli* falls in the low µM range.^30^ Thus, these initial reactions, performed at high enzyme concentration (12.5 µM, 50 mol% of tRNA^Pyl^), could mask reactivity differences that are relevant under conditions that better mimic the cellular environment. To more carefully characterize the relative reactivity of monomers **1**-**4**, we evaluated the yield of both mono- and di-acylated tRNA^Pyl^ as the concentration of *Ma*PylRS was reduced stepwise from 12.5 µM (50 mol%) to 625 nΜ (2.5 mol%) (Figure 2D). At low PylRS concentrations, the yield of tRNA^Pyl^ mono-acylated with *(S)*-β^2^-OH **3** was comparable to that of *(S)*-α-OH **2** and higher than that of (*S*)-α-NH_2_ **1** (Figure 2D); at higher enzyme concentrations, the di-acylated product predominates. The yield of tRNA^Pyl^ mono-acylated with the enantiomer *(R)*-β^2^-OH **4** is highest at high enzyme concentrations. These results imply that *(S)*-β^2^-OH **3** is a better substrate for *Ma*PylRS than its enantiomer *(R)*-β^2^-OH **4**, especially under conditions where enzyme concentration is limiting.

### *Ma*FRSA accepts β^2^-hydroxy acids as substrates *in vitro*

The PylRS derivative FRSA contains two active site mutations (N166A and V168A) that favor substrates with substituted Phe side chains.^23^ One of the best substrates for FRSA is the α-amino acid *m*-CF_3_-Phe **5 (**Figure 2E). To determine if *Ma*FRSA also accepts β^2^-HA substrates, we performed *in vitro* tRNA acylation reactions supplemented with *m*-CF_3_-Phe **5** alongside analogous reactions containing *(S)*-α-OH **6** and the enantiopure β^2^-OH analogs **7** and **8**. Reactions performed with 12.5 μM *Ma*FRSA, 25 μΜ *Ma*tRNA^Pyl^ and 10 mM **5** or **6** cleanly generated the expected mono-acylated tRNA products **5**-acyl-tRNA^Pyl^ (23175.6 Da) and **6**-acyl-tRNA^Pyl^ (23176.8 Da) (Figure 2E).

Quantification of the monoacyl-tRNA and unacylated tRNA pools indicates yields of 45% and 29% for **5**-acyl-tRNA^Pyl^ and **6**-acyl-tRNA^Pyl^ respectively (**Supplementary** Figure 3). Aminoacylation reactions supplemented with 10 mM *(S)*-β^2^-HA monomer **7** yielded only a single low signal peak in the TIC corresponding to the molecular weight of monoacyl-tRNA (23189.5 Da) with a yield of <1% **7**-acyl-tRNA^Pyl^ (**Supplementary** Figure 3). By contrast, supplementation of an analogous aminoacylation reaction with *(R)*-β^2^-HA **8** yielded two peaks in the TIC corresponding to monoacylated (23189.5 Da) and diacylated **8**-acyl-tRNA^Pyl^ (23420.1 Da) (Figure 2E) in 28% and 11% yield, respectively. Interestingly, while *Ma*PylRS processes *(S)*-β^2^-OH **3** more efficiently than *(R)*-β^2^-OH **4**, *Ma*FRSA shows the opposite preference, processing *(R)*-β^2^-HA **8** more efficiently than *(S)*-β^2^-HA **7.**

The analogous plot showing the yield of mono-+ di-acylated tRNA^Pyl^ as a function of [*Ma*PylRS] is shown in **Supplementary** Figure 2. (E) Deconvoluted mass spectra of *in vitro* tRNA^Pyl^ acylation reactions containing 12.5 µM *Ma*FRSA and supplemented with monomers **5-8**. Signal is normalized to the highest signal in respective traces. Expected masses shown correspond to mono-acylated products.

### *Ma*PylRS supports *in vivo* synthesis of a protein containing a single β^2^-HA

We next asked whether the *in vitro* tRNA^Pyl^ acylation efficiencies observed with β^2^-OH substrates would support the incorporation of these monomers into proteins biosynthesized in *E. coli*. Experiments were performed using the recoded *E. coli* strain C321.ΔΑ.exp, which lacks all endogenous TAG codons and release factor 1 (RF1).^31^ Cells were co-transformed with a pMega plasmid^25^ encoding the *Ma*PylRS/*Ma*tRNA^Pyl^ pair (pMega-*Ma*PylRS) as well as a pET22b plasmid encoding sfGFP with an in-frame TAG codon at position 3 (sfGFP-3TAG) (Figure 3A). *Ma*tRNA^Pyl^ naturally decodes TAG codons,^32^ making successful translation of full-length sfGFP is dependent on the concentration and activity of acyl-tRNA^Pyl^. Test expressions were performed in the presence of 0.5 to 2 mM **1**-**4** and both OD_600_ and 528 nm emission (F_528_) were monitored as a function of time (**Supplementary** Figure 4). Although the rate of increase in F_528_/OD_600_ was greater for monomers with a natural α-backbone (*(S)*-α-NH_2_ **1** and *(S)*-α-OH **2**), by 24 h all growth curves had reached saturation and hence this time point was used for comparisons of F_528_/OD_600_.

**Figure 3.**
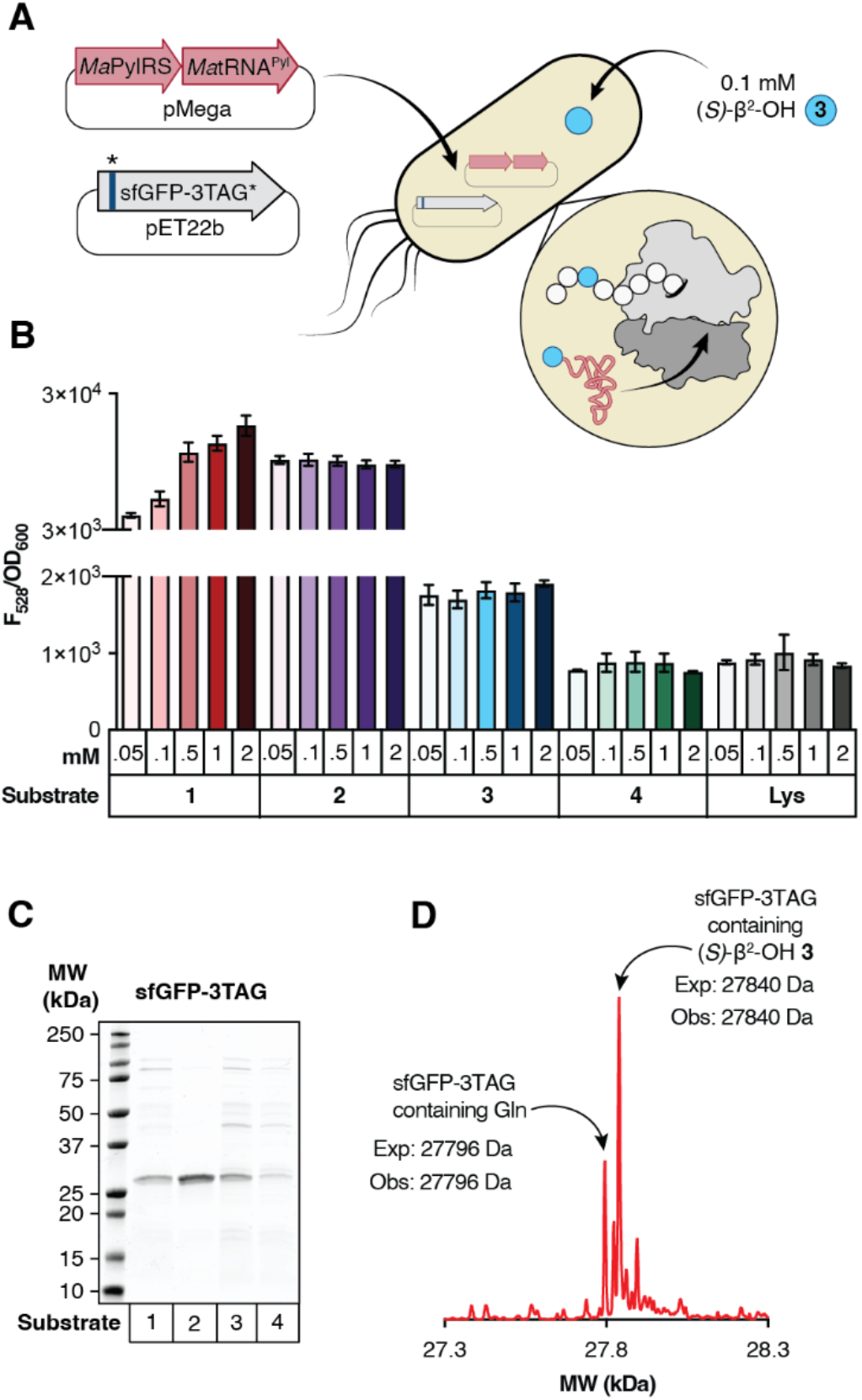
*Ma*PylRS supports *in vivo* synthesis of a protein containing a single β^2^-HA. (A) Workflow for protein expression in C321.ΔΑ.exp *E. coli* transformed with pMega-*Ma*PylRS and pET22b-sfGFP-3TAG. (B) Plot of F_528_/OD_600_ values measured 24 h after induction with 1 mM IPTG as a function of substrate identity and concentration. (C) SDS-PAGE of sfGFP-3TAG expressed in the presence of **1-4**. (D) Deconvoluted mass spectrum of sfGFP-3TAG expressed in the presence of (*S)*-β^2^-OH **3**.

Comparison of F_528_/OD_600_ values after 24 h revealed a clear concentration-dependent increase when cultures were supplemented with BocK **1** relative to those in which substrate was withheld (ΔAA) or supplemented with Lys, which is not a substrate for *Ma*PylRS (Figure 3B).^22^ Although no concentration dependence of the F_528_/OD_600_ value was observed when cultures were supplemented with *(S)*-α-OH BocK **2**, the F_528_/OD_600_ values observed after 24 h were comparable to those observed in the presence of 0.5 mM **1**. Cultures supplemented with *(S)*-β^2^-OH **3** also showed an increase in F_528_/OD_600_ relative to those in which substrate was withheld (ΔAA) or supplemented with Lys.

Interestingly, like growths supplemented with *(S)*-α-OH BocK **2**, the F_528_/OD_600_ values observed after 24 h, although low, were independent of the concentration of *(S)*-β^2^-OH **3** over an 80-fold range of concentration. No increases in F_528_/OD_600_ relative to background were observed when cultures were supplemented with *(R)*-β^2^-OH **4.** Identical protein expression assays with C321.ΔΑ.exp cells harboring *Ma*FRSA and supplemented with either enantiomer of β^2^-ΟΗ-*m*-CF_3_-Phe yielded no significant sfGFP expression over a ΔAA control (**Supplementary** Figure 5).

Two experiments were performed in an attempt to increase the F_528_/OD_600_ values of cells expressing sfGFP in the presence of *(S)*-β^2^-OH **3.** The growth experiments described above were repeated using either Top10 or BL21 (DE3) *E. coli* in place of C321.ΔΑ.exp. We also examined whether the F_528_/OD_600_ values of growths supplemented with *(S)*-β^2^-OH **3** could be improved via mutation of *Ma*tRNA^Pyl^. Previous work has emphasized the effect of tRNA identity on the efficiency of non-canonical α-amino acid incorporation into proteins.^33^ A rationally evolved, orthologous *Mb*tRNA^Pyl-opt^ bearing mutations at the base of the acceptor and T-stems improved the incorporation of certain non-canonical α-amino acid substrates into proteins expressed in BL21 (DE3) and Top10 E. coli,^23^ and mutations at the identical sites in an evolved *Ec*tRNA^Sec^ increased the incorporation of selenocysteine at TAG codons when grown in a derivative strain of C321.ΔA.^34^ Neither changing the expression strain nor the tRNA body improved the observed F_528_/OD_600_ values of growths supplemented with *(S)*-β^2^-OH **3** (**Supplementary** Figure 6).

To confirm that *(S)*-β^2^-OH **3** was being introduced into sfGFP-3TAG, we isolated sfGFP-3TAG from a preparative growth of C321.ΔΑ.exp cells transformed with pMega-*Ma*PylRS and pET22b-sfGFP-3TAG and supplemented with 0.1 mM *(S)*-β^2^-OH **3,** and characterized the product using SDS-PAGE (Figure 3C) and LC-HRMS (Figure 3D).

SDS-PAGE of cultures supplemented with (*S*)-β^2^-OH **3** produced a major protein product of ∼28 kDa whose mobility was comparable to proteins expressed in the presence of (*S*)-α-NH_2_ **1** and (*S*)-α-OH **2** (Figure 3C). The deconvoluted mass spectrum of purified sfGFP-3TAG expressed in the presence of *(S)*-β^2^-OH **3** included a major peak at 27841 Da, corresponding to the expected molecular mass of sfGFP with *(S)*-β^2^-OH **3** at position 3 but lacking residues 1-2 due to ester hydrolysis. A second, smaller peak at 27725 Da was also observed, corresponding to the mass of sfGFP with Gln at position 3; in this case residue 2 is retained. Although *(S)*-β^2^-OH **3** and *(R)*-β^2^-OH **4** are excellent substrates for PylRS *in vitro*, with activities comparable to that of (*S*)-α-NH_2_ **1**, only *(S)*-β^2^-OH **3** is incorporated into proteins in cells, and with lower efficiency than anticipated based only on aaRS activity *in vitro (vide infra)*.

To evaluate if a protein containing an intact β^2^-ester could be isolated, we designed three additional sfGFP expression plasmids in which a TAG codon was inserted in place of or between residues E213 and K214. These two residues can function as the N- and C-termini of a split sfGFP variant that assembles from two independent polypeptides.^35^ We reasoned that these sites would be well-suited to accommodate an internal β-ester without disrupting either the sfGFP fold or chromophore maturation. C321.ΔΑ.exp cells were co-transformed with pMega-*Ma*PylRS and a pET22b plasmid encoding sfGFP with an in-frame TAG codon at position 213 (pET22b-sfGFP-213TAG), position 214 (pET22b-214TAG), or between positions 213 and 214 (pET22b-213-TAG-214). Growths were supplemented with 0.1 mM **2**, **3**, or **4** as described previously, and both F_528_ and OD_600_ were monitored as a function of time (**Supplementary** Figure 7). Again, although the rate of increase in F_528_ was greater for growths containing *(S)*-α-OH BocK **2** than those containing *(S)*-β^2^-OH **3** or *(R)*-β^2^-OH **4**, by 24 h all growth curves had reached saturation and this time point was used for comparisons of F_528_/OD_600_. We observed a robust increase in the F_528_/OD_600_ signal of growths expressing sfGFP-213TAG, sfGFP-214TAG, or sfGFP-213-TAG-214 in the presence of *(S)*-α-OH **2** and a modest increase in F_528_/OD_600_ when cultures expressing sfGFP-213-TAG-214 were supplemented with *(S)*-β^2^-OH **3** (Figure 4A).

**Figure 4.**
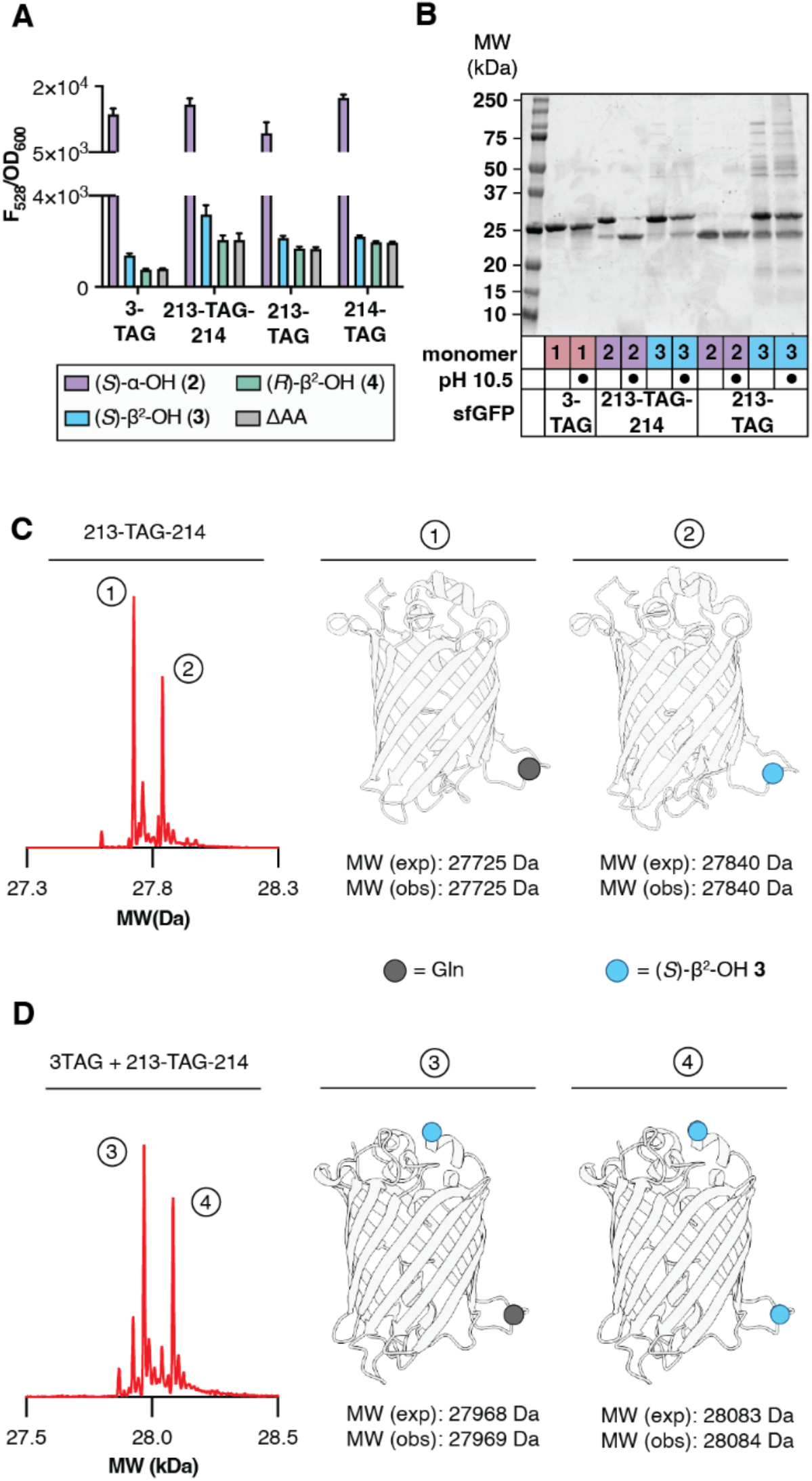
*Ma*PylRS supports the *in vivo* synthesis of sfGFP containing an intact β^2^-ester and two β^2^-hydroxy acids. (A) Plot of F_528_/OD_600_ values of C321.ΔΑ.exp *E. coli* transformed with pMega-*Ma*PylRS and the indicated sfGFP expression plasmid measured 24 h after induction with 0.1 mM IPTG as a function of monomer identity. (B) SDS-PAGE of the indicated isolated sfGFP variants after treatment with CAPS buffer (pH 10.5) or MilliQ for 2 h at 37 °C. (C) Deconvoluted mass spectrum of sfGFP-213-TAG-214 expressed in the presence of (*S)*-β^2^-OH **3** reveals two products. One contains a single residue of **3**, the other a single residue of Gln. (D) Deconvoluted mass spectrum of sfGFP-3-TAG-213-TAG-214 isolated from cells grown in defined media lacking Gln and the presence of 0.1 mM (*S)*-β^2^-OH **3.** Two products are observed whose masses correspond to (1) sfGFP with the addition of two copies of (*S)*-β^2^-OH **3** and (2) sfGFP with the addition of one copy of (*S)*-β^2^-OH **3** and a single Gln residue (**Supplementary** Figure 9).

sfGFP variants were isolated from preparative growths of C321.ΔΑ.exp cells transformed with pMega-*Ma*PylRS and either sfGFP-3TAG, sfGFP-213TAG, or sfGFP-213-TAG-214 and supplemented with BocK **1**, *(S)*-α-OH **2,** or *(S)*-β^2^-OH **3.** The products were characterized by SDS-PAGE, with and without base treatment to confirm the presence of an ester bond (pH 10.5) (Figure 4B), as well as by LC-HRMS (Figure 4C). SDS-PAGE analysis of sfGFP-213-TAG-214 isolated from C321.ΔΑ.exp cultures supplemented with *(S)*-α-ΟΗ **2** and without base treatment shows two bands, one that migrates with the MW expected for intact sfGFP (∼28 kDa) and one that migrates with the MW expected for the ester hydrolysis product (∼24 kDa). SDS-PAGE analysis of an analogous sample after base treatment led to virtually complete loss of the 28 kDa band and an increase in the intensity of the 24 kDa band, consistent with the presence of a base-labile ester bond. SDS-PAGE analysis of sfGFP-213-TAG-214 grown in the presence of (S)-β^2^-OH **3** also yielded two bands at ∼24 and 28 kDa prior to base treatment. Base treatment led to partial loss of the 28 kDa band and an increase in intensity of the 24 kDa band, suggesting that a fraction of the sample contained a base-labile ester linkage. Only a truncated product was isolated from growths programmed with sfGFP-213TAG and supplemented with *(S)*-α-OH **2**; in the case of an analogous growth supplemented with *(S)*-β^2^-OH **3**, both full length and truncated protein was observed, and the ratio was unaffected by base treatment. This observation implies that an ester linkage within sfGFP-213TAG is more hydrolytically labile than an ester linkage within sfGFP-213-TAG-214.

The successful internal incorporation of *(S)*-β^2^-OH **3** into sfGFP-213-TAG-214 was confirmed by LC-HRMS. sfGFP isolated from growths programmed with sfGFP-213-TAG-214 and supplemented with 0.1 mM *(S)*-β^2^-OH **3** contained two sfGFP variants. One contained a single residue of **3**, the other a single residue of Gln (Figure 4C). As predicted by gel analysis, sfGFP isolated from growths programmed with sfGFP-213-TAG and supplemented with *(S)*-β^2^-OH **3** revealed the presence of sfGFP with Gln at position 213 as well as a pair of protein fragments whose approximate molecular weights (∼4 kDa and ∼23.6 kDa) correspond to those expected if sfGFP was severed at position 213. However, the exact mass of the large (23625 Da) fragment was 18 Da less than that predicted on the basis of sequence alone (**Supplementary** Figure 8A). It is possible that ester hydrolysis in this case is promoted by the Asn residue at position 212, which could induce ester cleavage via the pathway used by asparagine lyase self-cleaving enzymes, the product of which undergoes dehydration (**Supplementary** Figure 8B).^36^

### *Ma*PylRS supports the *in vivo* synthesis of sfGFP containing two β^2^-HA monomers

Having identified that *(S)*-β^2^-OH **3** could be introduced into sfGFP at position 3 as well as between positions 213 and 214, we next asked whether this monomer could be introduced at both positions simultaneously. C321.ΔA.exp cells were transformed with pMega-*Ma*PylRS as well as a plasmid encoding sfGFP with TAG codons at position 3 as well as between E213 and K214 (pET22b-sfGFP-3TAG-213-TAG-214), and grown in the presence of 0.1 mM *(S)*-β^2^-OH **3** or 0.1 mM *(S)*-α-OH **2**. LC-HRMS analysis of the isolated sfGFP generated in the presence of *(S)*-β^2^-OH **3** confirmed the introduction of this monomer at both positions, albeit with Gln contamination at one or both positions (**Supplementary** Figure 9). However, expression of sfGFP-3TAG-213-TAG-214 with 0.1 mM *(S)*-β^2^-OH **3** in defined media lacking Gln^25,37^ led to only two products. One contained *(S)*-β^2^-OH **3** at position 3 and glutamine at position 213-214 (27969 Da); the other contained *(S)*-β^2^-OH **3** at both positions (28083 Da) (Figure 4D).

### Metadynamics simulations probe enantioselectivity of the PTC with respect to β^2^-OH-monomers

For over thirty years, the level of non-canonical α-amino acid incorporation at a stop codon has been used as a proxy for aaRS activity *in vivo*.^38^ This proxy fails for the β^2^-backbone monomers studied here. Although *Ma*PylRS acylates tRNA^Pyl^ with both *(S)*-β^2^-OH **3** and *(R)*-β^2^-OH **4** at levels comparable to *(S)*-α-ΝΗ_2_ **1** and *(S)*-α-ΟH **2** *in vitro,* only *(S)*-β^2^-OH **3** is introduced into protein in cells. We turned to metadynamics to learn more about how *(S)*-α-NH_2_ **1,** *(S)*-α-OH **2**, *(S)*-β^2^-OH **3** and *(R)*-β^2^-OH **4** are accommodated within the ribosomal A-site when loaded on tRNA^Pyl^, as previously reported simulations emphasize the important relationship between nucleophile positioning within the PTC and bond formation.^39^ These metadynamics simulations made use of a reduced ribosome model (RRM) containing fMet-tRNA^fMet^ in the P site and either **1-**acyl-tRNA^fMet^, **2-**acyl-tRNA^fMet^, **3-**acyl-tRNA^fMet^, or **4-**acyl-tRNA^fMet^ in the A site. Two 100 ns metadynamics simulations were initiated using two distinct monomer poses and the results were averaged. One pose aligned the A-site nucleophile with the nucleophilic atom of the A-site Met in the 2.1 Å cryo-EM model used to build the RRM;^40^ this pose placed the α-NH_2,_ α-OH, or β^2^-OH nucleophile as close as possible to the P-site carbonyl. The second pose was generated by rotating the psi (ψ) angle of the β^2^-OH monomer by 180°; this pose placed the β^2^-OH nucleophile further away from the P-site carbonyl.

Previous results suggest that reactivity within the PTC is related to two distinct parameters: the N_α_-C_sp2_ distance between the A-site nucleophile (N_α_) and the P-site carbonyl electrophile (C*_sp2_*), and the Bürgi-Dunitz attack angle (α_BD_). Monomers that react readily within the PTC populate a conformational space characterized by a N_α_-C_sp2_ distance of <4 Å and a α_BD_ value between 76 and 115°.^40^ Examination of plots showing α_BD_ as a function of the N_α_–C_sp2_ distance reveal global minima that differentiate highly reactive (*(S)*-α-NH_2_ **1,** *(S)*-α-OH **2**), moderately reactive (*(S)*-β^2^-OH **3**), and non-reactive (*(R)*-β^2^-OH **4**) monomers (Figure 5A). The free energy surface for an RRM containing **1-**acyl-tRNA^fMet^ in the A-site is defined by an N_α_-C_sp2_ distance of 3.9 ± 0.2 Å and α_BD_ of 92.3° ± 13.5° averaged for all final poses within 1 kcal/mol of the global energy minimum (Figure 5B, darkest blue). Both of these values are comparable to those reported for an RRM with Met-tRNA^fMet^ in the A-site (N_α_-C_sp2_ = 3.7 Å and α_BD_ = 76°).^40^ Simulations with **2-**acyl-tRNA^fMet^ in the A-site yielded comparable values (N_α_-C_sp2_ = 3.9 ± 0.1 Å and α_BD_ = 69.8 ± 3.6°), which is fully consistent with the high reactivity of *(S)*-α-OH **2** *in vivo*.

**Figure 5.**
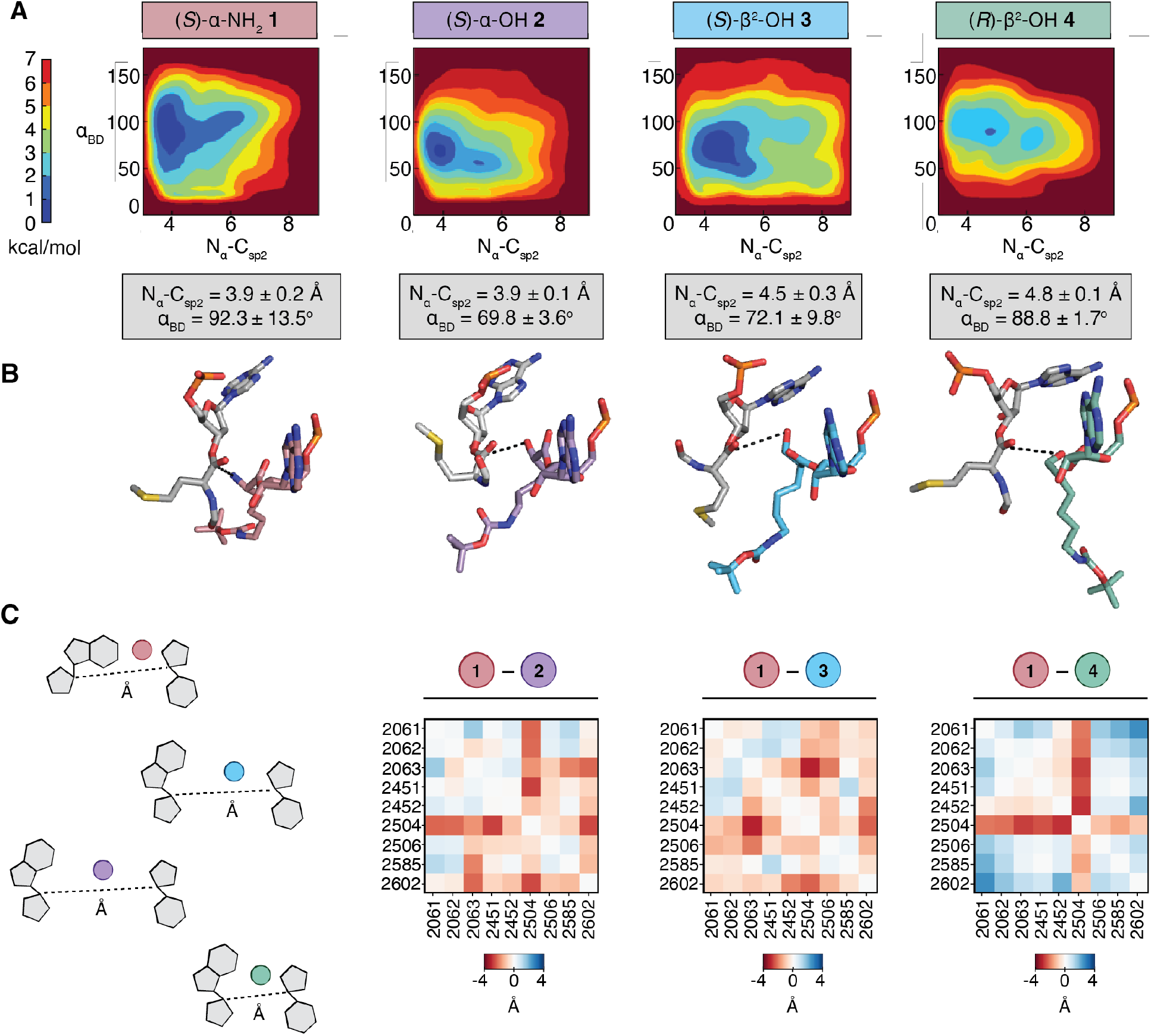
Metadynamics simulations of β^2^-OH-BocK prior to ester bond formation in a Reduced Ribosome Model (RRM). (A) Free Energy Surface (FES) plots of 30 Å RRM containing a P-site tRNA^Met^ acylated with fMet and A-site tRNA^Met^ acylated with monomers **1**–**4**, plotted along the collective variables of Bürgi–Dunitz angle *α*_BD_ and N_α_–C*_sp_*_2_ distance. Each FES shown is the average of two metadynamics runs starting from orientations of the A-site monomers that differ by a 180° rotation about the ψ angle. The color scale represents the free energy in kilocalories per mole (kcal/mol), where the global minima are set at 0 and therefore the various heights of the energy scales are based on the energetics of the fluctuations of the A-site monomers. Average N_α_-C_sp2_ distances and α_BD_ angles for the 0-1 kcal/mol energy minima of each plot are displayed below (gray boxes) (B) A representative conformation and relative geometry of the P-site (gray) and A-site (colored) monomers from the frames at the free energy minimum of each FES plot. Poses were chosen to highlight the N_α_–C*_sp_*_2_ distance (black dotted line) and the Bürgi–Dunitz angle *α*_BD_ at the free energy minimum. The tRNAs to which each monomer is attached as well as the RRM surrounding the two monomers have been omitted for clarity. (C) PIA plots displaying the pairwise-distance differences between the C1^’^ carbons of nucleotides within 5 Å of A-site monomers along the metadynamics trajectories. Heatmaps were constructed by subtracting the C1^’^ distances for respective monomers from C1^’^ distances for *(S)*-α-NH_2_ **1**. Darker red indicates greater distances between nucleotides for the respective monomer relative to monomer **1**, darker blue indicates smaller distances between nucleotides for respective monomers relative to monomer **1**.

The free energy surfaces for an RRM containing **3-**acyl-tRNA^fMet^ or **4-**acyl-tRNA^fMet^ are defined by different global minima values. For the RRM containing **3-**acyl-tRNA^fMet^ in the A-site, we observe low energy poses characterized by N_α_-C_sp2_ distances of 4.5 ± 0.3 Å and α_BD_ values of 72.1° ± 9.8°. The N_α_-C_sp2_ distance for **3-**acyl-tRNA^fMet^ falls outside the N_α_-C_sp2_ distance range for monomers predicted to be highly reactive in the ribosome and suggests that the relatively low incorporation of *(S)*-β^2^-OH **3** is due in part to poor sampling of conformations within the PTC that support rapid bond formation. The free energy surface of an RRM containing **4-**acyl-tRNA^fMet^ is defined by a similar averaged N_α_-C_sp2_ distance and α_BD_ but with much smaller standard deviations (N_α_-C_sp2_ = 4.8 ± 0.1 Å and α_BD_ = 88.8° ± 1.7°). These differences suggest that **3-**acyl-tRNA^fMet^ can achieve N_α_-C_sp2_ distances that permit modest incorporation of *(S)*-β^2^-OH **3** into a ribosomal product whereas **4-**acyl-tRNA^fMet^ cannot. Indeed, the energy minima of metadynamics trajectories for monomer **4** produces a minimized structure in which the β-OH of **4-**acyl-tRNA^fMet^ is turned away from the P-site carbonyl electrophile (Figure 5B). This analysis provides one plausible explanation for the observed *in vivo* differences in reactivity of A-site tRNAs esterified with *(S)*-β^2^-OH **3** and *(R)*-β^2^-OH **4**.

We next examined the overall structure of the PTC during each trajectory to learn more about conformational changes in the ribosome that might facilitate productive incorporation of *(S)*-β^2^-OH **3** and not *(R)*-β^2^-OH **4**. We first calculated the average distance between the C1’ atoms of all rRNA bases within 5 Å of either acyl-tRNA^fMet^ in the P- or A-site of RRMs containing **1-**acyl-tRNA^fMet^, **2-**acyl-tRNA^fMet^, **3-**acyl-tRNA^fMet^, or **4-**acyl-tRNA^fMet^ (Figure 5C). The matrix of pairwise interactions in each of the four minimized RRMs provides an effective map of how the PTC responds to structurally and stereochemically distinct ⍺-NH_2_, ⍺-OH, and β^2^-OH monomers. To visualize differences between the maps, the pairwise interaction distances calculated from RRMs containing **2-**acyl-tRNA^fMet^, **3-**acyl-tRNA^fMet^, or **4-**acyl-tRNA^fMet^ in the A-site were subtracted from the pairwise interactions distances calculated from the RRM containing **1-**acyl-tRNA^fMet^. The resulting pairwise interaction difference analysis (PIA) plots provide a comprehensive view of how the internal architecture of the PTC varies in a monomer-dependent fashion (Figure 5C).^41,42^ In a PIA plot, distances between rRNA bases that are greater in the presence of **2**, **3-** or **4**-acyl-tRNA^fMet^ relative to **1**-acyl-tRNA^fMet^ are represented in red, whereas distances that are shorter are represented in blue.

Examination of the PIA plot for RRMs containing **1-**acyl-tRNA^fMet^ or **2**-acyl-tRNA^fMet^ shows significant differences at only two rRNA bases (U2504, located within H89, and C2063, part of the highly conserved A2450-C2063 non-Watson-Crick base pair). In both cases the differences reflect an expansion of the PTC when bound to **2**-acyl-tRNA^fMet^.

The PIA plot for RRMs containing **1-**acyl-tRNA^fMet^ or **3**-acyl-tRNA^fMet^ is highly similar to the plot comparing the RRMs for **1-**acyl-tRNA^fMet^ and **2**-acyl-tRNA^fMet^ with the largest differences again involving lengthened interactions with A2504 and C2063.

In contrast, the PIA plot comparing the RRMs for **1-**acyl-tRNA^fMet^ and **4-**acyl-tRNA^fMet^ is different. First, it highlights many pairwise interactions that are markedly shorter when **4-** acyl-tRNA^fMet^ occupies the A-site, including those involving U2506, U2585, and A2062. Changes involving U2506 and U2585 are especially notable, as both have been implicated as critical for induced conformational changes required for efficient bond formation.^43^ U2585 is believed to shield the P-site peptidyl-tRNA from hydrolysis in the uninduced state and rotate away in the induced state to expose the ester bond for nucleophilic attack by the A-site monomer.^43^ Indeed, recent cryo-EM structures of *E. coli* ribosomes with aminobenzoic acid monomers in the A-site show U2585 locked in the uninduced conformation, prohibiting access to the P-site peptidyl-tRNA.^44^ Overall, the decreased pairwise distances within the RRM containing **4-**acyl-tRNA^fMet^ may indicate an inability to support dynamic movements necessary for efficient bond formation. These differences in RRM structure provide a second explanation for the observed *in vivo* enantioselective preference for *(S)*-β^2^-OH **3** over *(R)*-β^2^-OH **4.** More broadly, they emphasize that the complex mechanism of translation, including (but not limited to) tRNA-induced conformational changes and essential bound water molecules^44^ must be considered in future ribosome or monomer engineering efforts.

## Discussion

The programmed synthesis of sequence-defined biomaterials whose monomer backbones diverge from canonical α-amino acids represents the next frontier in protein and biomaterial evolution. Such next-generation molecules provide otherwise non-existent opportunities to develop improved biological therapeutics, bioremediation tools, and biodegradable plastic-like materials. But progress towards this goal has been exceptionally slow. Although most elements of the translational machinery tolerate even wildly divergent ⍺-amino acid side chains,^38^ altered backbones are largely tolerated only *in vitro*,^45^ at small scale, under non-physiological conditions, and with efficiencies that have not been rigorously evaluated.

Here we report that β^2^-hydroxy acids possessing both *(R)* and *(S)* absolute configuration are excellent substrates for pyrrolysyl-tRNA synthetase (PylRS) enzymes *in vitro* and that *(S)*-β^2^-hydroxy acid are substrates *in cellulo*. One unexpected finding is that the *(S)*-β^2^-hydroxy acid incorporated successfully into protein by the ribosome *in vivo* possesses an absolute configuration that maps onto a D-α-amino acid, not an L-α-amino acid. Although the structure of the PylRS variant FRSA bound to 2-benzylmalonate^25^ provides an explanation for the absence of enantio-preference at the aaRS level, the enantio-preference for the D-like species in the PTC was a surprise.

Metadynamics simulations and recent cryo-EM structures^44^ provide evidence that this enantio-preference has less to do with inherent stereochemistry than with differences in the ability to induce conformational changes within the PTC necessary for an induced state fit and favorable bond formation. A better understanding of the interactions necessary to facilitate P-site electrophile deshielding could be used to identify D-α-amino acids that are successfully elongated *in vivo*, another long-sought goal.^46^

Overall, the combined biochemical and computational approach reported here provides the first example of orthogonal cellular translation through a β-backbone and an important steppingstone towards cell-based synthesis of β^2^-HA/α-AA hybrids with new-to-nature functionalities. They also improve our understanding of how the WT *E. coli* ribosome accommodates substrates with non-native stereochemistry and backbone configurations and suggest that translation factor^47^ and/or ribosomal engineering,^20,48,49^ as well as alternative approaches,^50^ may be needed to achieve robust levels of β-linkages within proteins produced in cells.

## Author contributions

Study conception and design: N.X.H., A.S.; preparation of materials: N.X.H.; data collection: N.X.H., A.M.A. analysis and interpretation of results: N.X.H., A.M.A., A.S.; and manuscript preparation: N.X.H., A.S.

## Acknowledgements

We are grateful to members of the Schepartz, Chatterjee, and Cate labs for insightful comments and suggestions. We also thank Dr. Chandrima Majumdar for productive discussions about induced state ribosomal conformations. This work was supported by the NSF Center for Genetically Encoded Materials (C-GEM; CHE 2002182). A.S. is a Chan-Zuckerberg Biohub-San Francisco Investigator and an ARC Innovation Investigator.

## TOC Graphic

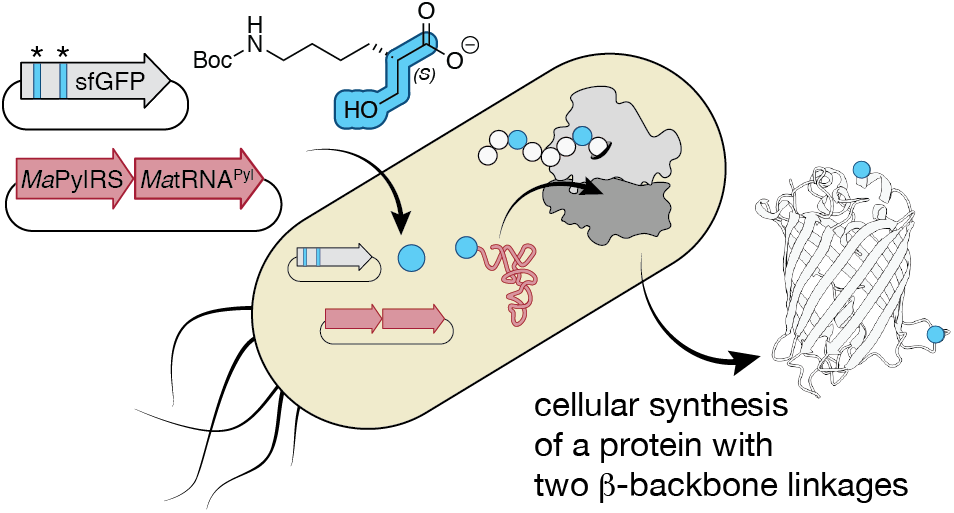

