## Supplementary Information for "Incorporation of multiple β^2^-backbones into a protein *in vivo* using an orthogonal aminoacyl-tRNA synthetase"

### Table of Contents

I. Materials and general information

  A. General Cloning Protocols

II. aaRS expression and purification

  A. MaPylRS expression plasmid and coding sequence

  B. MaPylRS expression and purification

  C. MaFRSA expression plasmid and coding sequence

  D. MaFRSA expression and purification

III. Synthesis of MatRNA$^{Pyl}$

  A. In vitro synthesis of dsDNA encoding *Ma*tRNA$^{Pyl}$

  B. In vitro transcription of *Ma*tRNA$^{Pyl}$

  C. Purification and characterization of *Ma*tRNA$^{Pyl}$

IV. in vitro tRNA acylation reactions

  A. Procedure for acylation of *Ma*tRNA$^{Pyl}$ in vitro

  B. LC-MS analysis of *Ma*tRNA$^{Pyl}$

V. Cloning of sfGFP-TAG variants

  A. Design of sfGFP-TAG variants

  B. Cloning sfGFP variants into pET22b vector

  C. Expression of sfGFP variants

    1. Plate reader-based expression assay of sfGFP variants

    2. Preparative scale expression of sfGFP variants

    3. Preparative scale expression of sfGFP variants in defined media

    4. Base hydrolysis and SDS-PAGE of sfGFP with internal β$^{2}$-esters

VI. Metadynamics simulations of β$^{2}$-hydroxy acid-tRNA in the ribosome

VII. Supplementary Figures 1-11

VIII. Supplementary Tables 1 -

  A. Table 1: DNA oligos used

  B. Table 2: Recipe for defined Δgln media

IX. Supplementary References

### I. General information

##### A. General Cloning Protocols

Antibiotics used in this study were supplied at concentrations recommended by addgene.org for selection of E. coli harboring a plasmid of interest (carbenicillin = 100 μg/mL, spectinomycin = 50 μg/mL)

##### B. General heat shock transformation Protocol

All cell strains used in this study (XL1 blue, Top10, DH5α, BL21, C321.ΔA.exp) were chemically competent and transformed with desired plasmids according to the following heat shock transformation protocol. A 100 μL stock of chemically competent cells was thawed on ice for 10 min, after which either 1 μL of 100-200 ng/μL purified plasmid or 5 μL of KLD-treated plasmid was added to cells on ice and incubated for 30 min. Cells were then subjected to heat shock at 45$^o$C for 30 s and then placed back on ice for 5 min. After, 900 μL of SOC Outgrowth Medium (NEB, cat # B9020S) was added to heat shocked cells. 100 μL of cells transformed with a plasmid harboring resistance to carbenicillin were plated directly on LB-agar with carbencillin with no recovery step. Plasmids for all other antibiotic resistances were allowed to recover at 37$^o$C with shaking at 200 rpm for 1 h. After, 100μL of recovered cells were plated on a pre-warmed LB-agar + carbenicillin plate and left to grow overnight at 37$^o$C in a dry air incubator.

##### C. Plasmid outgrowth and purification from glycerol stocks

For plasmids available as glycerol stocks stored at -80$^o$C in the lab, the following general outgrowth and purification protocol was used. A toothpick stab of DH5α or XL1-blue cells harboring the relevant plasmid was streaked onto an LB-agar plate containing the requisite antibiotic for plasmid selection and grown overnight at 37$^o$C in a dry air incubator. The following day, 2 individual colonies were picked and used to inoculate a starter culture consisting of 5 mL LB media + antibiotic. This starter culture was grown overnight in a shaking incubator at 37$^o$C and 200 rpm. The following day, overnight cultures were pooled and plasmid was purified using a QIAprep® Spin Miniprep Kit (cat# 27104) following the manufacturer's protocol. Purified plasmids were stored at -20$^{o}$C.

##### 1. Site-directed mutagenesis for plasmid variants

sfGFP and *Ma*tRNA$^{Pyl}$ variants were synthesized by site-directed mutagenesis using the KLD Enzyme Mix (NEB, Cat #M0554S) per the following manufacturer's reaction protocol. Briefly, 1 µL of a 2 ng/µL stock of a parent plasmid added to 12.5 µL of Q5® Hot Start High-Fidelity DNA Polymerase along with 1.25 µL of 10 µM forward and reverse primers (**Supplementary Table 2**) designed for site-directed mutagenesis using NEBaseChanger (nebasechanger.neb.com). The reaction mixture was brought up to 25 µL with MilliQ $H_2O$ and subjected to around-the-horn PCR to generate a linearized dsDNA plasmid harboring the desired mutations. 1 µL of around-the-horn PCR product was combined with 1 µL of 10x KLD Enzyme Mix, 5 µL of 2x KLD Reaction Buffer, and 3 µL of MilliQ $H_2O$. The reaction mixture was mixed well by pipetting up and down 5 times and then incubated at room temperature for 5 min. 5 µL of the KLD-treated reaction was then transformed into chemically competent XL1 blue cells according to the **General Transformation Protocol** above.

#### II. aaRS expression and purification

##### A. MaPylRS expression plasmid and coding sequence

The plasmid used to express *Ma*PylRS (pET32A-MaPylRS) was obtained from DH5α glycerol stocks previously published.[1] This plasmid encodes the complete sequence of **Pyrrolysyl-tRNA synthetase** from *Methanomethylophilus alvus* (Uniprot ID: M9SC49) preceded by an N-terminal **GSS-6xHis-SSG** sequence to enable immobilized metal ion affinity chromatography (IMAC) purification.

M**GSSHHHHHHSSG**LVPRGSH**MTVKYTDAQIQRLREYGNGTYEQKVFEDLASRDAAF**
**SKEMSVASTDNEKKIKGMIANPSRHGLTQLMNDIADALVAEGFIEVRTPIFISKDALAR**
**MTITEDKPLFKQVFWIDEKRALRPMLAPNLYSVMRDLRDHTDGPVKIFEMGSCFRKE**

**SHSGMHLEEFTMLNLVDMGPRGDATEVLKNYISVVMKAAGLPDYDLVQEESDVYKETIDVEINGQEVCSAAVGPHYLDAAHDVHEPWSGAGFGLERLLTIREKYSTVKKGGASISYLNGAKIN***

##### B. MaPylRS expression and purification

1 µL of 100-200 ng/µL of purified pET32A-MaPylRS plasmid was transformed into chemically competent *E. coli* BL21(DE3) cells (NEB) according to the manufacturer's protocol and plated onto LB-agar plates containing 100 µg/mL carbenicillin. Plates were incubated overnight at 37°C in a dry air incubator. The following day a single colony of *E. coli* BL21(DE3) carrying pET32A-*Ma*PylRS was used to inoculate a 5 mL starter culture of LB media + 100 µg/mL carbenicillin and grown overnight at 37°C with shaking at 200 rpm. The following day, 1 mL of starter culture was used to inoculate 100 mL of terrific broth (TB) with 100 µg/mL carbenicillin and grown at 37°C with shaking at 200 rpm. When cultures reached $OD_{600}$ = ~1.0, protein expression was induced by addition of IPTG to 1 mM final concentration and incubated at 18°C with shaking at 200 rpm for 18 h. The following day, cells were harvested via centrifugation in a Beckman Coulter Allegra® X-14R Benchtop Centrifuge in a Beckman SX4750 Swinging Bucket Rotor at 4300 x g for 1 h. Clarified media was decanted and cell pellets were resuspended in 10 mL Lysis Buffer (50 mM $K_2HPO_4$, 25 mM imidazole, 500 mM NaCl, 5 mM 2-mercaptoethanol, pH 7.4) + 1 cOmplete™, Mini, EDTA-free Protease Inhibitor Cocktail tablet (Roche). The cell suspension was passed through at 18 gauge needle to break up cell clumps and cells were lysed via homogenization (Avestin Emulsiflex C3). Briefly, the homogenizer was primed with 50 mL Lysis Buffer after which cell suspension was added and outlet tubing was placed above the inlet cup creating a closed loop. Cells were then passed through the homogenizer at a flow rate of 16.7 mL/min and lysed via homogenization with 15,000-20,000 PSI pulses for 2 min. After homogenization, lysate was collected into a fresh 50 mL conical tube. Lysate was clarified by centrifugation in a Beckman Coulter Allegra® X-14R Benchtop Centrifuge in a FX6100 Fixed-Angle Aluminum Rotor- 6 x 100 mL at 11,000 x g for 1 h. During lysate clarification, 2 mL (1 mL of packed resin) of TALON® resin slurry (Takara Bio) was equilibrated into lysis buffer by first spinning down resin at 1000 x g for 1 min and decanting storage solution

following by 4 subsequent resuspensions and spins in lysis buffer. TALON® resin was added to clarified lysate in a fresh 50 mL conical tube and allowed to equilibrate via batch binding at 4ºC on a rotisserie for 1 hr. After, lysate-resin slurry was poured into a disposable Poly-Prep® Chromatography Column (Bio-Rad, cat # 7311550) and flow through was collected. Bound resin was washed with 30 column-volumes (30 mL) of Lysis Buffer and collected as 3 x 10 mL washes. Bound protein was then eluted using 2.5 mL Elution Buffer (50 mM potassium phosphate, 500 mM imidazole, 500 mM sodium chloride, 5 mM 2-mercaptoethanol, pH 7.4). Eluate was desalted using a PD-10 desalting columns packed with Sephadex G-25 resin (Cytiva, cat # 17085101) using manufacturer's instructions and eluted in 3.5 mL Storage Buffer (100 mM NaCl, 100 mM HEPES, 10 mM $MgCl_2$, 4 mM DTT, 20% v/v glycerol, pH 7.2). Protein was concentrated to a final concentration of 1 mM using Amicon Ultra-2 Centrifugal Filter Unit MWCO 10 kDa (Millipore Sigma, cat # C7715) and divided into 10 µL aliquots. Aliquots were flash-frozen in liquid $N_2$, stored at -80ºC, and used as single-use aliquots.

##### C. MaFRSA expression plasmid and coding sequence

The plasmid used to express MaFRSA (pET32A-MaFRSA) was obtained from DH5α glycerol stocks previously published.[1] This plasmid encodes the complete sequence of **FRSA**, an engineered variant of PylRS carrying to mutations in the active site (**N166A** & **V168A**) that alter enzyme preference for substrates with ring-substituted phenylalanine sidechains,[2] preceded by an N-terminal **GSS-6xHis-SSG** sequence to enable immobilized metal affinity chromatography (IMAC) purification.

M**GSSHHHHHHSSG**LVPRGSH**MTVKYTDAQIQRLREYGNGTYEQKVFEDLASRDAAF**
**SKEMSVASTDNEKKIKGMIANPSRHGLTQLMNDIADALVAEGFIEVRTPIFISKDALAR**
**MTITEDKPLFKQVFWIDEKRALRPMLAPNLYSVMRDLRDHTDGPVKIFEMGSCFRKE**
**SHSGMHLEEFTMLALADMGPRGDATEVLKNYISVVMKAAGLPDYDLVQEESDVYKE**
**TIDVEINGQEVCSAAVGPHYLDAAHDVHEPWSGAGFGLERLLTIREKYSTVKKGGASI**
**SYLNGAKIN***

##### D. MaFRSA expression and purification

Expression and purification of *Ma*FRSA followed a protocol that was identical to that employed for the purification of *Ma*PylRS (section IIB).

#### III. Synthesis of *Ma*tRNA$^{Pyl}$

##### A. *In vitro* synthesis of dsDNA encoding *Ma*tRNA$^{Pyl}$

A double stranded DNA template encoding for *Ma*tRNA$^{Pyl}$ was synthesized from complementary DNA oligonucleotides (*Ma*PylT-F and *Ma*PylT-R, oligos 1-2, **Supplementary Table 1**) ordered from IDT (Supplementary Table X). Briefly, each oligo resuspended to a stock concentration of 100 mM and 1 µL was added to a PCR tube and diluted to a final concentration of 2 mM upon addition of 23 µL of MilliQ $H_2O$ and 25 µL of GoTaq® G2 Master Mix (Promega). Oligos were annealed and extended using the following protocol on a Bio-Rad C1000 Touch Thermal Cycler: 94 °C for 30 s, 30 cycles of 94 °C for 20 s, 53 °C for 30 s and 68 °C for 60 s, and finally 68 °C for 300 s.

After the annealing and extension protocol, reactions were supplemented with sodium acetate (pH 5.2) to a final concentration of 300 mM and washed 1x with 25:24:1 (v/v/v) phenol:chloroform:isoamyl alcohol. The aqueous layer was washed 2x with chloroform and the dsDNA product was precipitated by addition of ice cold 200-proof ethanol to a final concentration of 71% (v/v). Samples were placed in a dry ice acetone bath for 30 min and pelleted by centrifugation at 213000 x g, 4$^{o}$C for 30 min. Liquid was decanted and pellets were washed once in 71% ethanol followed by a second centrifugation. dsDNA pellets were then resuspended in 100 µL MilliQ $H_2O$, quantified using a NanoDrop ND-1000 Spectrophotometer and diluted to a final concentration of 500 ng/µL. The final pure dsDNA template has a **T7 promoter** immediately preceded by a **C** to increase T7 transcript yield and a **2’-methoxy modification** on the penultimate guanosine of the reverse complement strand to reduce non-templated addition of ribonucleotides by T7 RNA polymerase.

**DNA sequence for *Ma*PylT:**

5'-

**C****TAATACGACTCACTATA**GGGGGACGGTCCGGCGACCAGCGGGTCTCTAAAACC

TAGCCAGCGGGGTTCGACGCCCCGGTCTCTCGCC**m**A-3'

**Full length transcription product for *Ma*tRNA$^{Pyl}$:**

5'-

GGGGGACGGUCCGGCGACCAGCGGGUCUCUAAAACCUAGCCAGCGGGGUUCGA

CGCCCCGGUCUCUCGCCA-3'

**B. *In vitro* transcription of *Ma*tRNA$^{Pyl}$**

*Ma*-tRNA$^{Pyl}$ was transcribed in vitro using a modified version of a published procedure.[3] Transcription reactions (25 µL) contained the following components: 40 mM Tris-HCl (pH 8.0), 100 mM NaCl, 20 mM DTT, 2 mM spermidine, 5 mM ATP, 5 mM cytidine triphosphate, 5 mM guanosine triphosphate, 5 mM uridine triphosphate, 20 mM guanosine monophosphate, 0.2 mg/mL bovine serum albumin, 20 mM $MgCl_2$, 12.5 ng/µLDNA template and 0.025 mg/mL T7 RNA polymerase. The reaction mixtures were incubated at 37 °C in a thermocycler for 3 h. To each 25 µL reaction was added 3.125 U of RQ1 RNAse-free DNAse I (Promega) and 3.125 µL of 10x RQ1 DNAse buffer. Reactions were then incubated at 30$^{o}$C for 30 min. After incubation, 8x transcription reactions were pooled (250 µL total) and sodium acetate (pH 5.2) was added to a final concentration in 300 µL. The transcription reaction mixtures were then extracted once with a 1:1 (v/v) mixture of acidic phenol (pH 4.5) and chloroform and washed twice with chloroform. To the samples was added ice cold 200 proof ethanol to a final concentration of 71% (v/v) and incubated for 30 min in a dry ice-acetone bath. Samples were spun at 21300 x g for 30 min at 4$^{o}$C to pellet RNA, after which liquid was decanted and RNA was dried for 20 min. To remove small molecules, the tRNA was resuspended in MilliQ $H_2O$ and further purified using Bio-Rad Micro Bio-Spin™ P-30 Gel Columns, Tris Buffer (RNase-free) after first exchanging the column buffer with MilliQ $H_2O$ according to the manufacturer's protocol. The tRNA was precipitated once more in ice cold 71% ethanol, resuspended in water, quantified using a NanoDrop ND-1000 Spectrophotometer, aliquoted and stored at −20 °C.

##### C. Purification and characterization of *Ma*tRNA$^{Pyl}$

tRNA was analyzed by urea–PAGE (**Supplementary Figure 10A**) using Bio-Rad 10% Mini-PROTEAN® TBE-Urea Precast Gel . Gels were run at 120 V for 30 min and then stained with SYBR Safe Gel Stain (Thermo Fisher) for 5 min before imaging. *Ma*-tRNA$^{Pyl}$ was also analyzed by LC–MS to confirm its identity. Samples were resolved on an ACQUITY UPLC BEH C18 column (130 Å, 1.7 μm, 2.1 mm × 50 mm, 60 °C; Waters, 186002350) using an ACQUITY UPLC I-Class PLUS instrument (Waters, 186015082). The mobile phases used were 8 mM triethylamine, 80 mM hexafluoroisopropanol and 5 μM EDTA (free acid) in 100% MilliQ $H_2O$ (mobile phase A) and 4 mM triethylamine, 40 mM hexafluoroisopropanol and 5 μM EDTA (free acid) in 50% MilliQ water–50% methanol (mobile phase B). The analysis was performed at a flow rate of 0.3 ml min$^{-1}$ and began with mobile phase B at 22%, increasing linearly to 40% B over 10 min, followed by a linear gradient from 40% to 60% B for 1 min, a hold at 60% B for 1 min, a linear gradient from 60% to 22% B over 0.1 min and then a hold at 22% B for 2.9 min. The mass of the RNA was analyzed by LC–MS with a Xevo G2-XS Tof instrument (Waters, 186010532) in negative ion mode with the following parameters: capillary voltage = 2,000 V, sampling cone = 40, source off-set = 40, source temperature = 140 °C, desolvation temperature = 20 °C, cone gas flow = 10 L/h, desolvation gas flow = 800 L/h and collection rate = 1 spectrum/s. The expected masses of the oligonucleotide products were calculated using the AAT Bioquest RNA Molecular Weight Calculator. Deconvoluted mass spectra were obtained using the MaxEnt software (Waters). A representative mass spectra of *Ma*tRNA$^{Pyl}$ is included in **Supplementary Figure 10B**.

#### IV. *in vitro* tRNA acylation reactions

##### A. Procedure for acylation of *Ma*tRNA$^{Pyl}$ *in vitro*

The reaction mixtures (25 μL) used to acylate *Ma*tRNA$^{Pyl}$ contained the following components: 100 mM HEPES-K (pH 7.5), 4 mM DTT, 10 mM $MgCl_2$, 10 mM ATP, 10 mM enantiopure substrate (**1 - 4,** *Ma*PylRS; **5 - 8** *Ma*FRSA), 0.1 U Pyrophosphatase, Inorganic (*E. coli*) (NEB), 25 μM *Ma*tRNA$^{Pyl}$ and 0-12.5 μM synthetase (either *Ma*PylRS or *Ma*FRSA). The reaction mixtures were incubated at 37 °C in a dry air incubator for

2 h. After, sodium acetate (pH 5.2) was added to the acylation reactions to a final concentration of 300 mM in a volume of 200 µL. The reaction mixtures were then extracted once with a 1:1 (v/v) mixture of acidic phenol (pH 4.5) and chloroform and washed twice with chloroform. After extraction, the acylated tRNA was precipitated by adding ethanol to a final concentration of 71% and incubation in a dry ice-acetone bath for 30 min, followed by centrifugation at 21300 x *g* for 30 min at 4 °C. After carefully aspirating supernatant, tRNA pellets were left to dry for 15 min at room temperature. Pellets were then resuspended in 2.0 µL of RNAse-free MilliQ $H_2O$ 1 µL of which was diluted 1:20 in RNAse-free MilliQ $H_2O$ and transferred to a high recovery mass spec vial for LC-MS analysis.

##### B. LC-MS analysis of *Ma*tRNA$^{Pyl}$

The tRNA samples from the enzymatic acylation reactions were analyzed by LC–MS as described in **Purification and characterization of *Ma*tRNA$^{Pyl}$** (Supplementary Information Section VIII, B.). Because the unacylated tRNA peak in each TIC contained tRNA species that could not be enzymatically acylated (primarily tRNAs that lack the 3′-terminal adenosine),[4] simple integration of the acylated and non-acylated peaks in the absorbance at 260 nm ($A_{260}$) chromatogram could not accurately quantify the acylation yield. To accurately quantify the acylation yield, we used a published procedure.[1,5] For each sample, mass data were collected between *m/z* = 500 and 2,000. A subset of the mass data collected defined as the raw MS deconvolution range was used to produce the deconvoluted mass spectra. The raw MS deconvolution range of each macromolecule species contained multiple peaks corresponding to different charge states of that macromolecule. Within the raw mass spectrum deconvolution range we identified the most abundant charge state peak in the raw mass spectrum of each tRNA species (unacylated, monoacylated and diacylated tRNA). To quantify the relative abundance of each species, the exact mass of the major ions (±0.3000 Da) was extracted from the TIC to produce the EICs. The EICs were integrated and the areas of the peaks that aligned with the correct peaks in the TIC (as determined from the deconvoluted mass spectrum) were used to quantify the yields. The expected masses of the oligonucleotide products were calculated using the AAT Bioquest RNA Molecular

Weight Calculator, and the molecular masses of the small molecules added to them were calculated using ChemDraw 19.0.

### V. Cloning of *Ma*tRNA$^{Pyl-opt}$ variants

#### A. Design of *Ma*tRNA$^{Pyl-opt}$ variants

pMega-*Ma*PylRS-*Ma*tRNA$^{Pyl}$ was purified from a glycerol stock of DH5α cells with using the plasmid outgrowth and purification protocol described in **General Cloning Protocols** with the use of spectinomycin as the antibiotic selector.

We designed 2 additional pMega-*Ma*PylRS plasmids bearing either *Ma*tRNA$^{Pyl-opt1}$ or *Ma*tRNA$^{Pyl-opt2}$, chimeric tRNAs with mutations originating from evolved *Mb*tRNA$^{Pyl-opt}$ or *Ec*tRNA$^{Sec}$ respectively (**Supplementary Figure 6**) and synthesized them via site-directed mutagenesis.

#### B. Cloning *Ma*tRNA$^{Pyl-opt}$ variants into pMega-*Ma*PylRS vector

pMega-*Ma*PylRS plasmids encoding either *Ma*tRNA$^{Pyl-opt1}$ or *Ma*tRNA$^{Pyl-opt2}$were synthesized by site-directed mutagenesis as described under **Site-directed mutagenesis protocol** using pMega-*Ma*PylRS-*Ma*tRNA$^{Pyl}$ as the parent plasmid. 5 µL of the KLD-treated reaction was then transformed into chemically competent XL1-Blue cells as described under **General heat shock transformation protocol**. After heat shock transformation and recovery, 100µL of recovered cells were plated on a pre-warmed LB-agar + spectinomycin plate and left to grow overnight at 37$^{o}$C in a dry air incubator. After overnight growth, 5 individual colonies per construct from overnight growths were picked and used to inoculate 5mL of LB + spectinomycin and grown overnight in a shaking incubator at 200 rpm and 37$^{o}$C. Plasmid from liquid cultures was isolated and purified using a QIAprep® Spin Miniprep Kit (cat# 27104) per manufacturer's instructions using a vacuum manifold. Typical yields were in the range of 45 µL of 150-300 ng/µL of pure plasmid. Correct variants were sequence verified by sanger sequencing using the Berkeley DNA Sequencing Core Facility and whole plasmid sequencing from Primordium Labs.

### VI. Cloning of sfGFP-TAG variants

#### A. Design of sfGFP-TAG variants

pET22b-sfGFP-3TAG was purified from a glycerol stock of DH5α cells with an identical outgrowth and plasmid purification protocol as for pET32A-*Ma*FRSA and pET32A-*Ma*PylRS.

We designed 3 additional sfGFP-TAG variants for β-hydroxy acid incorporation with consideration for sites that would not disrupt protein folding or chromophore maturation. We hypothesized that mutating residues E213 or K214 to an amber codon, or inserting an amber codon between them, would achieve this criteria as "cutting" between these residues forms a functional split sfGFP[6] as well as a stable, fluorescent circular permutation.[7]

* = amber codon

**sfGFP-E213TAG protein sequence:**

MSKGEELFTGVVPILVELDGDVNGHKFSVRGEGEGDATNGKLTLKFICTTGKLPVPWP
TLVTTLTYGVQCFSRYPDHMKRHDFFKSAMPEGYVQERTISFKDDGTYKTRAEVKFE
GDTLVNRIELKGIDFKEDGNILGHKLEYNFNSHNVYITADKQKNGIKANFKIRHNVEDGS
VQLADHYQQNTPIGDGPVLLPDNHYLSTQSVLSKDPN*KRDHMVLLEFVTAAGITHGM
DELYKGSHHHHHH

**sfGFP-E213TAGK214 protein sequence:**

MSKGEELFTGVVPILVELDGDVNGHKFSVRGEGEGDATNGKLTLKFICTTGKLPVPWP
TLVTTLTYGVQCFSRYPDHMKRHDFFKSAMPEGYVQERTISFKDDGTYKTRAEVKFE
GDTLVNRIELKGIDFKEDGNILGHKLEYNFNSHNVYITADKQKNGIKANFKIRHNVEDGS
VQLADHYQQNTPIGDGPVLLPDNHYLSTQSVLSKDPNE*KRDHMVLLEFVTAAGITHG
MDELYKGSHHHHHH

**sfGFP-K214TAG protein sequence:**
MSKGEELFTGVVPILVELDGDVNGHKFSVRGEGEGDATNGKLTLKFICTTGKLPVPWP
TLVTTLTYGVQCFSRYPDHMKRHDFFKSAMPEGYVQERTISFKDDGTYKTRAEVKFE
GDTLVNRIELKGIDFKEDGNILGHKLEYNFNSHNVYITADKQKNGIKANFKIRHNVEDGS
VQLADHYQQNTPIGDGPVLLPDNHYLSTQSVLSKDPNE*RDHMVLLEFVTAAGITHGM
DELYKGSHHHHHH

#### B. Cloning sfGFP variants into pET22b vector

pET22b plasmids carrying sfGFP variants with an in frame amber (TAG) codon were synthesized by site-directed mutagenesis as described under **Site-directed mutagenesis protocol** using pET22b-sfGFP3TAG as the parent plasmid. 5 µL of the KLD-treated reaction was then transformed into chemically competent XL1-Blue cells as described under **General heat shock transformation protocol**. The pET22b plasmid encodes resistance to carbenicillin so after heatshock and resuspension cells were immediately plated on LB-agar + carbenicillin and left to grow overnight at 37$^{o}$C in a dry air incubator. After overnight growth, 5 individual colonies per construct from overnight growths were picked and used to inoculate 5mL of LB + carbenicillin and grown overnight in a shaking incubator at 200 rpm and 37$^{o}$C. Plasmid from liquid cultures was isolated and purified using a QIAprep® Spin Miniprep Kit (cat# 27104) per manufacturer's instructions using a vacuum manifold. Typical yields were in the range of 45 µL of 100-200 ng/µL of pure plasmid. Correct sfGFP variants were sequence verified by sanger sequencing using the Berkeley DNA Sequencing Core Facility and whole plasmid sequencing from Primordium Labs.

#### C. Expression of sfGFP variants

##### 1. Plate reader-based expression assay of sfGFP variants

Chemically competent *E. coli* cells (C321.ΔA.exp, BL21 (DE3), or Top10) were doubly transformed with (1) synthetase plasmid (pMega-*Ma*PylRS or pMega-*Ma*FRSA) and (2) sfGFP reported plasmid (pET22b-sfGFP-3TAG, pET22b-sfGFP-213TAG, pET22b-sfGFP-213-insTAG-214, or pET22b-sfGFP-214TAG) and plated onto selective LB-Agar (100µg/mL carbenicillin + 50µg/mL spectinomycin). The following day, a single colony per double transformant was picked and used to inoculate 5mLs of LB + carb + spec, and grown overnight at 37°C with shaking at 200 rpm. The following day, 200 µL of overnight culture was used to inoculate 19.8 mL of TB + carb + spec, and grown at 37°C with shaking at 200 rpm until reaching an $OD_{600}$ = ~1.0 - 1.2 (typically 4.5 - 5 h). To 646.75 µL of cell culture at appropriate $OD_{600}$ was added IPTG (1 mM final concentration) and monomer (10 mM final concentration). To a Corning® 96-well Flat Clear Bottom Black Polystyrene TC-treated Microplate (Cat# 3904), 200 µL of cell culture + IPTG and monomer was added per well with n = 3 technical replicates per condition. The plate was sealed with a gas permeable Breathe-Easy® sealing membrane (USA Scientific, Cat # 9123-6100). The plate was incubated at 37 °C for 24 h with continuous shaking in an Agilent Synergy HTX Multi-Mode Plate Reader. Two readings were made at 10 min intervals: (1) the absorbance at 600 nm, to measure cell density, and (2) sfGFP fluorescence with excitation at 485 nm and emission at 528 nm.

#### 2. Preparative scale expression of sfGFP variants

pMega-*Ma*PylRS and one pET22b-sfGFP expression plasmid (3TAG, 213TAG, 213-TAG-214, or 214-TAG)were transformed into C321.ΔA.exp as described in **Plate reader-based expression assay of sfGFP variants**. From transformants plated on selective LB-agar a single colony was picked and used to inoculate 5 mLs of LB + carb + spec and grown overnight at 37°C with shaking at 200 rpm. The following day, 1 mL of overnight starter culture was added to 99 mL of liquid TB + carb + spec and monomer **1**, **2**, **3**, or **4** was added to a final concentration of 0.1 mM. Growths were incubated at 37°C with shaking at 200 rpm until reaching an $OD_{600}$ = ~1.0 - 1.2 and expression of *Ma*PylRS and sfGFP was induced by addition of IPTG (1 mM final concentration). Growths were incubated at 37°C with shaking at 200 rpm for 18 h.

The following day, cells were harvested via centrifugation in a Beckman Coulter Allegra® X-14R Benchtop Centrifuge in a Beckman SX4750 Swinging Bucket Rotor at 4300 x g for 1 h. Clarified media was decanted and cell pellets were resuspended in 10 mL Lysis Buffer (50 mM $NaH_2PO_4$, 300 mM NaCl, pH 6.8) + 1 cOmplete™, Mini, EDTA-free Protease Inhibitor Cocktail tablet (Roche). The cell suspension was passed through at 18 gauge needle to break up cell clumps and cells were lysed via homogenization (Avestin Emulsiflex C3). Briefly, the homogenizer was primed with 50 mL Lysis Buffer after which cell suspension was added and outlet tubing was placed above the inlet cup creating a closed loop. Cells were then passed through the homogenizer at a flow rate of 16.7 mL/min and lysed via homogenization with 15,000-20,000 PSI pulses for 2 min. After homogenization, lysate was collected into a fresh 50 mL conical tube. Lysate was clarified by centrifugation in a Beckman Coulter Allegra® X-14R Benchtop Centrifuge in a FX6100 Fixed-Angle Aluminum Rotor- 6 x 100 mL at 11,000 x g for 1 h. During lysate clarification, 1 mL (0.5 mL of packed resin) of TALON® resin slurry (Takara Bio) was equilibrated into Lysis Buffer by first spinning down resin at 1000 x g for 1 min and decanting storage solution following by 4 subsequent resuspensions and spins in Lysis Buffer. TALON® resin was added to clarified lysate in a fresh 50 mL conical tube and allowed to equilibrate via batch binding at 4°C on a rotisserie for 1 hr. After, lysate-resin slurry was poured into a disposable Poly-Prep® Chromatography Column (Bio-Rad, cat # 7311550) and flow through was collected. Bound resin was washed with 30 column-volumes (30 mL) of Lysis Buffer and collected as 3 x 10 mL washes. Bound protein was then eluted using 2.5 mL Elution Buffer (50 mM $NaH_2PO_4$, 250 mM Imidazole, pH 6.8). Eluate was desalted using a PD-10 desalting columns packed with Sephadex G-25 resin (Cytiva, cat # 17085101) using manufacturer's instructions and eluted in 3.5 mL Storage Buffer (50 mM $NaH_2PO_4$, 250 mM NaCl, pH 6.8). Protein was quantified using a NanoDrop ND-1000 Spectrophotometer and concentrated using Amicon Ultra-2 Centrifugal Filter Unit MWCO 10 kDa as necessary for analysis by SDS-PAGE and LC-HRMS.

#### 3. Preparative scale expression of sfGFP variants in defined media

sfGFP variants expressed in a defined media followed the exact same expression and purification protocols as the above section apart from media formulation. The minimal media recipe was adapted from a published protocol.[8,9] See **Supplementary Table 2** for components of the defined media used in this study.

#### 4. Base hydrolysis and SDS-PAGE of sfGFP with internal $\beta^2$-esters

15 µL of a 10 µM stock of purified protein was denatured at 95°C for 5 min in a Bio-Rad C1000 Touch Thermal Cycler. 6 µL of 500 mM CAPS buffer (pH 10.5) was added to denatured protein and total sample volume was brought up to 30 µL with MilliQ $H_2O$. To negative controls was added 15 µL of MilliQ $H_2O$. Samples were then incubated at 37°C for 2 h to facilitate base hydrolysis. After, 6 µL of 5x loading dye (5% β-mercatoethanol, 0.02% bromophenol blue, 30% glycerol, 10% SDS, 250 mM Tris pH 6.8). 30 µL of sample was loaded per well to a Bio-Rad Any kD™ Mini-PROTEAN® TGX™ Precast Protein Gel and run at 200 V for 30 min. Gels were incubated in coomassie brilliant blue staining solution for 30 min followed by destaining solution for 30 min and imaged on a Bio-Rad ChemiDoc™ MP Imaging System.

#### 5. LC-HRMS Analysis of sfGFP

Proteins were analyzed by LC–MS to confirm their identity. The samples analyzed by MS were resolved using a Poroshell StableBond 300 C8 column (2.1 mm × 75 mm, 5 µm; Agilent, 660750-906) with a 1290 Infinity II ultra-high-performance liquid chromatograph (UHPLC; Agilent, G7120AR). The mobile phases used for separation were 0.1% formic acid in water (mobile phase A) and 100% acetonitrile (mobile phase B), and the flow rate was 0.4 ml $min^{-1}$. After an initial hold at 5% B for 0.5 min, the proteins were eluted using a linear gradient from 5% to 75% B for 9.5 min, a linear gradient from 75% to 100% B for 1 min, a hold at 100% B for 1 min, a linear gradient from 100% to 5% B for 3.5 min and finally a hold at 5% B for 4.5 min. The protein masses were analyzed by LC–HRMS using a 6530 Q-TOF AJS-ESI (Agilent, G6530BAR) instrument. The following parameters were used: gas temperature = 300 °C, drying gas flow = 12 l $min^{-1}$, nebulizer pressure = 35 psi, sheath gas temperature = 350 °C, sheath gas flow = 11 l $min^{-1}$, fragmentor voltage = 175 V, skimmer voltage = 65 V, peak-to-peak voltage ($V_{pp}$) = 750 V, capillary voltage ($V_{cap}$) = 3,500 V, nozzle voltage = 1,000 V and collection rate = 3 spectra $s^{-1}$.

#### VII. Metadynamics simulations of $\beta^2$-hydroxy acid-tRNA in the ribosome

As described in the Results, the starting point for our MD simulations was the RRM as reported in our previous study. The structure was solvated using the simple point charge water model.[10] K+ and Cl– ions corresponding to 0.15 M concentration were added as well as K+ counterions to neutralize the system. The final simulation box measured 95 Å along each side and consisted of ~88,000 atoms. The OPLS4 force field[11] and Desmond MD system (Schrödinger Release 2022-2) as implemented within Schrödinger Suite (release 2023-2) were used in this study. For all the non-α-amino acid monomers, the Force Field Builder (Schrödinger release 2023-2)[11] was used to parametrize the missing torsions. The systems were initially minimized and equilibrated with restraints on all solute heavy atoms, followed by production runs with all but the outer 10 Å C1′ and Cα atoms unrestrained. The constant-temperature, constant-pressure (NPT; number of particles *N*, pressure *P*, temperature *T*) ensemble was used with constant temperature at 300 K and Langevin dynamics. Desmond[12] (Schrödinger release 2023-2) was used for the metadynamics runs. The metadynamics production runs were carried out in duplicate (starting from different conformations as outlined above) for 100 ns each. The Nα–C*sp*2 distance and the Bürgi–Dunitz angle were used as collective variables. The biasing Gaussian potential ('hill') of 0.01 kcal mol–1 was used, and a width of 0.15 Å for the Nα–C*sp*2 distance and 2.5° for the Bürgi–Dunitz angle *α*BD were applied. Analysis of the runs was performed with Schrödinger's Python API (Schrödinger release 2023-2) as well as in-house Python scripts.

### VIII. Supplementary Figures 1-11

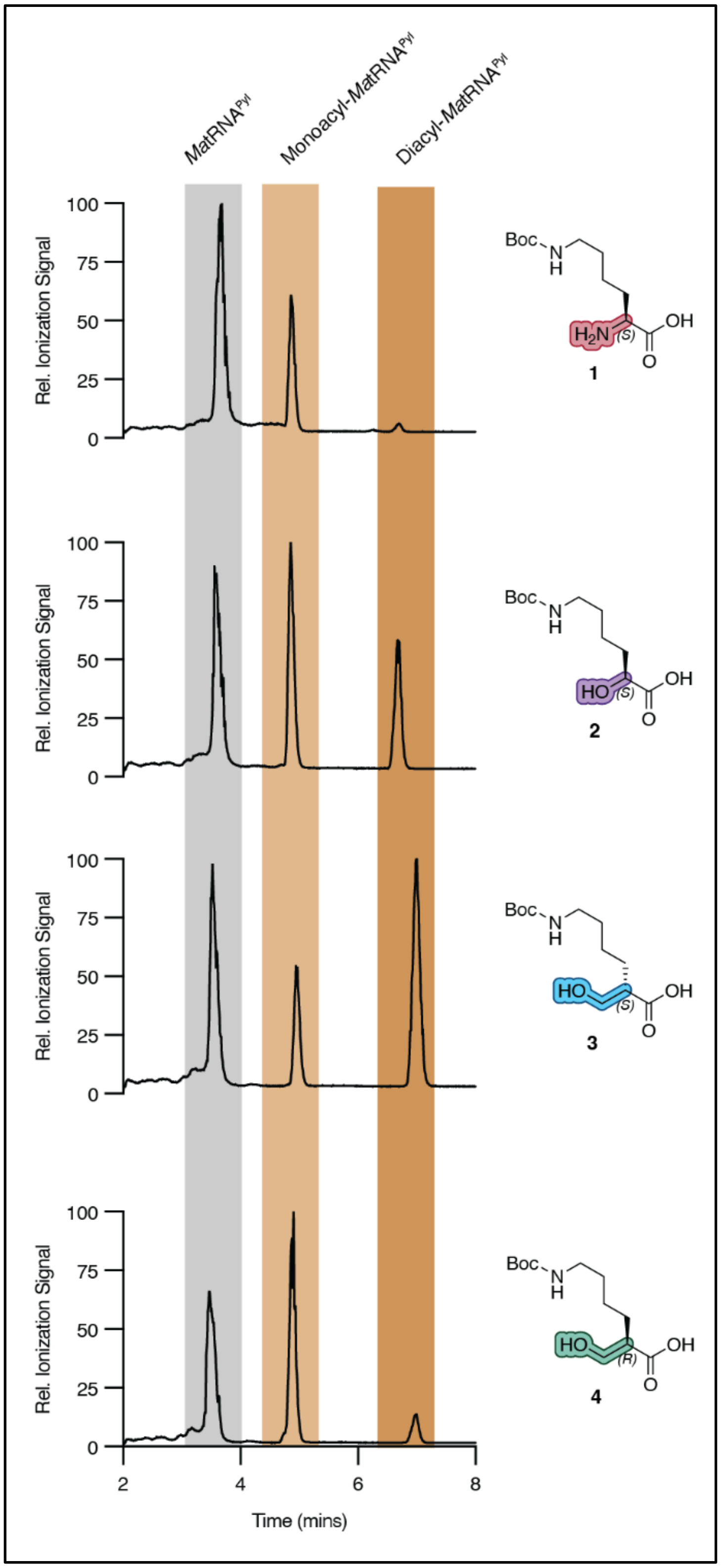

**Supplementary Figure 1.** ***Ma*****PylRS accept $\beta^2$-hydroxy acids as substrates *in vitro.*** Shown are representative traces from intact tRNA LC-HRMS analysis of reactions containing 12.5 µM *Ma*PylRS, 25 µM *Ma*tRNA$^{Pyl}$, and 25 µM of substrates **1-4.** Yields were calculated according to the workflow described in **Supplementary Information Section IV, B.**

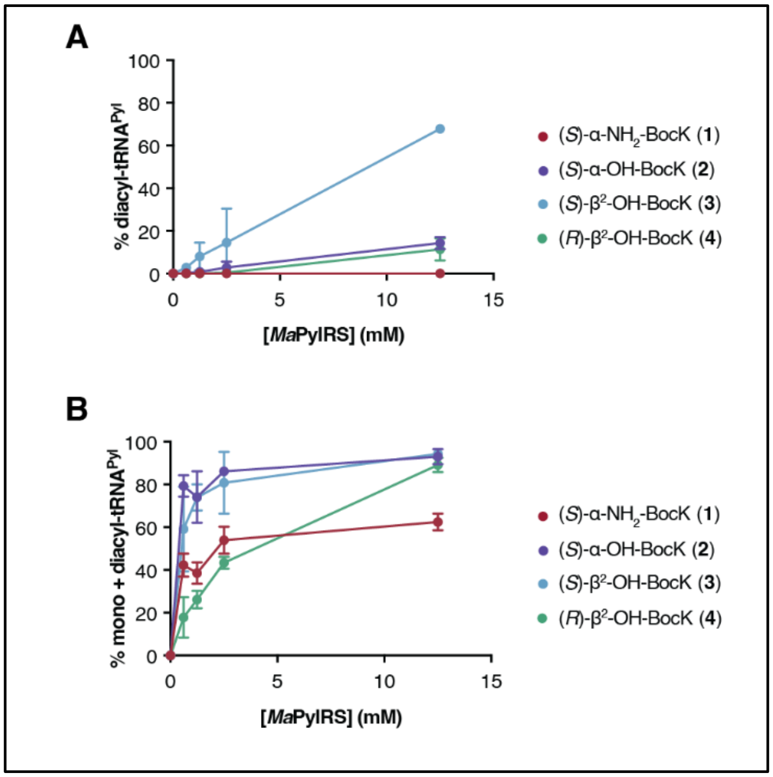

**Supplementary Figure 2. Effect of [*Ma*PylRS] on the fractional yields of diacyl-tRNA$^{Pyl}$ and total (monoacyl + diacyl)-tRNA$^{Pyl}$.** (A) Plot of data from intact tRNA LC-HRMS illustrating the % diacyl-tRNA$^{Pyl}$ observed in reactions containing substrates **1**-**4** as a function of [*Ma*PylRS]. n = 2, technical replicates. (B) Plot of data from intact tRNA LC-HRMS illustrating the % acyl-tRNA$^{Pyl}$ (mono + diacyl) observed in reactions containing substrates **1**-**4** as a function of [*Ma*PylRS]. n = 2, technical replicates.

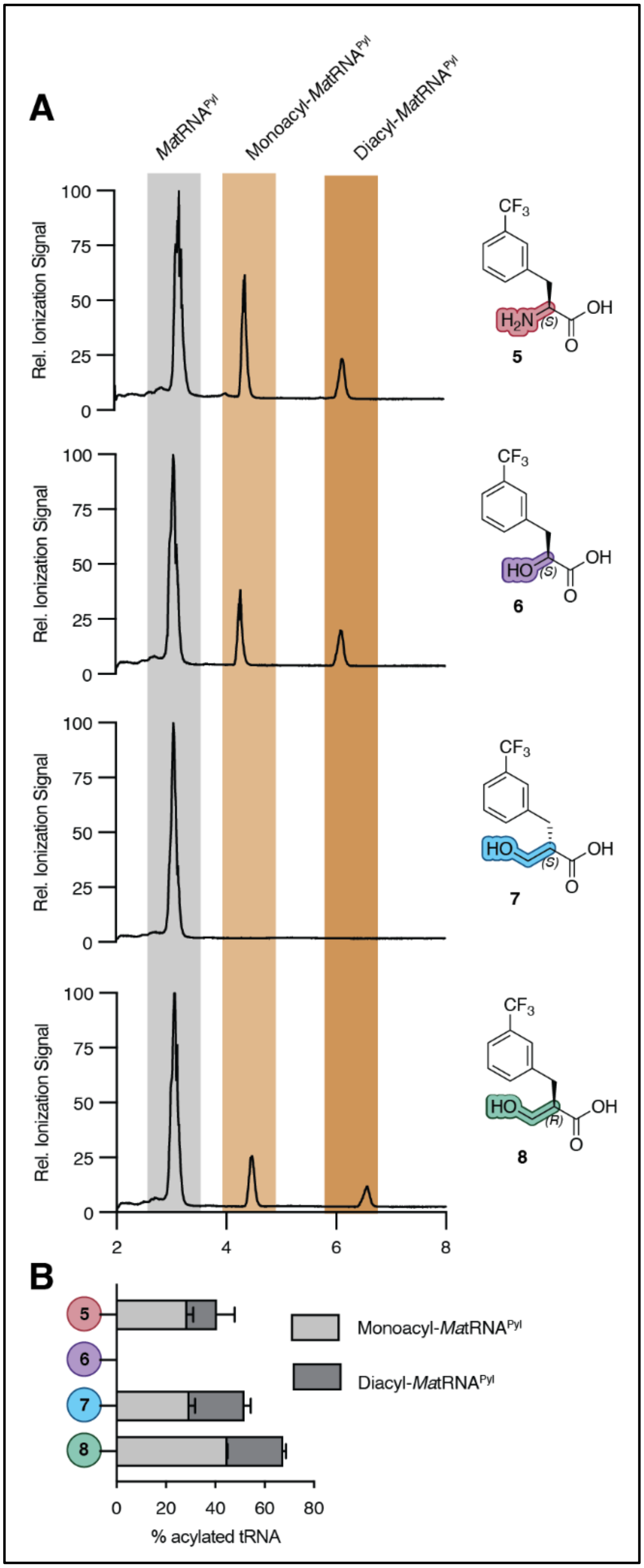

**Supplementary Figure 3.** ***Ma*FRSA accepts β²-hydroxy acids as substrates *in vitro.*** (A) Shown are representative traces from intact tRNA LC-HRMS analysis of reactions containing 12.5 μM *Ma*FRSA, 25 μM *Ma*tRNA$^{Pyl}$, and 25 μM of substrates **5-8.** After 2 h, these reactions show evidence of residual tRNA$^{Pyl}$ as well as mono- and diacylated tRNA products. Yields were calculated as described previously.[13] (B) Plot of data from intact tRNA LC-HRMS illustrating the relative fraction of monoacyl- and diacyl-tRNA$^{Pyl}$ in reactions supplemented with monomers **5**-**8**. n = 2, technical replicates.

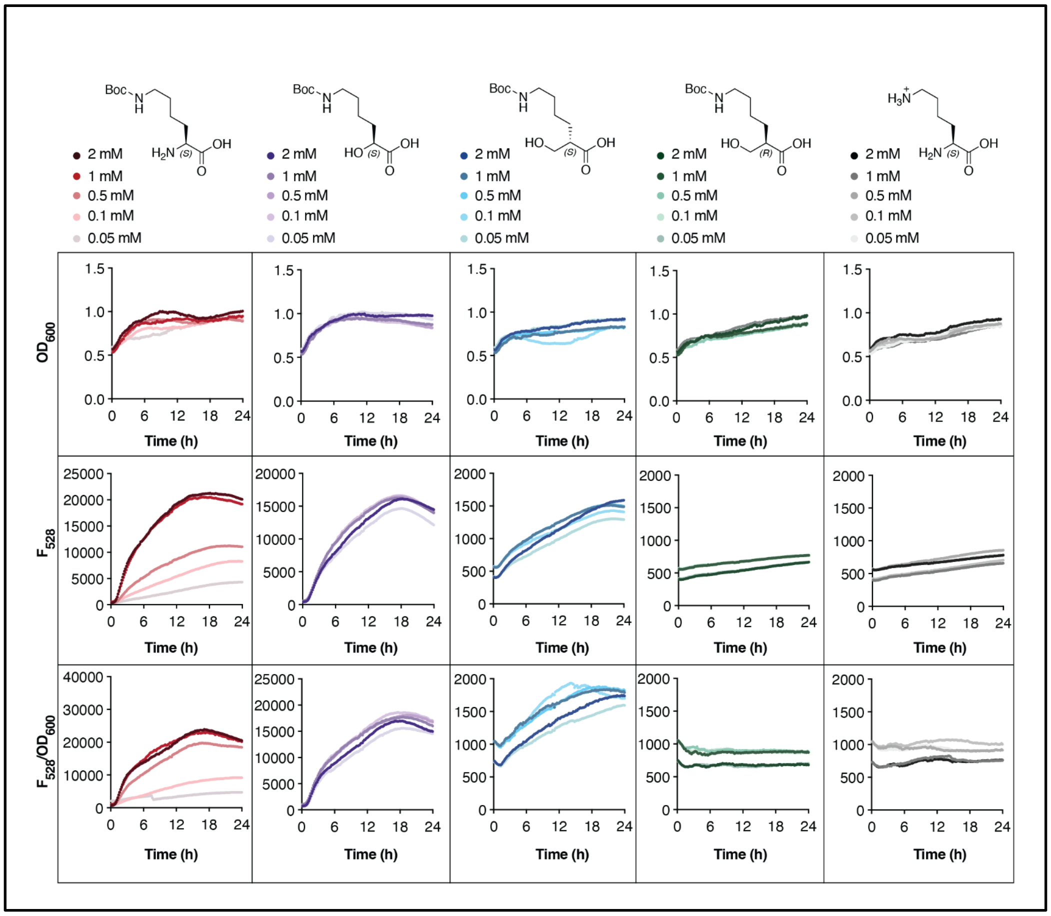

**Supplementary Figure 4. Time-dependent changes in cell density ($OD_{600}$) and 528 nm emission ($F_{528}$) of C321.ΔA.exp *E. coli* harboring pMega-*Ma*PylRS and pET22b-sfGFP-3TAG and grown in the presence of the indicated potential substrates.** Shown are growth ($OD_{600}$), GFP fluorescence ($F_{528}$), and growth-corrected ($F_{528}/OD_{600}$) curves for C321.ΔA.exp cells supplemented with varying concentrations (0.05 mM - 2 mM) of substrates **1** - **4**. Growth and expression conditions used are reported in **Supplementary Information Section V: Plate reader-based expression assay of sfGFP variants**.

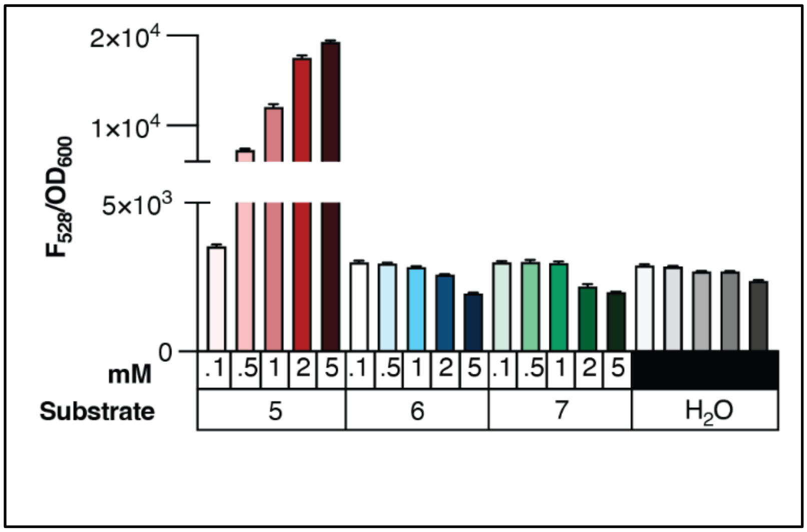

**Supplementary Figure 5. *Ma*FRSA does not support the incorporation of β$^2$-OH-*m*-$CF_3$-Phe into sfGFP-3TAG.** Shown is the $F_{528}/OD_{600}$ signal of C321.ΔA.exp *E. coli* harboring pMega-*Ma*FRSA and pET22b-sfGFP-3TAG and supplemented with the indicated substrates at 24 h post-induction with 1 mM IPTG. Although growths containing α-$NH_2$-*m*-$CF_3$-Phe **5** show a concentration-dependent increase in $F_{528}/OD_{600}$, those supplemented with *(S)*-β$^2$-OH **7** and *(R)*-β$^2$-OH **8** show no increase in $F_{528}/OD_{600}$ relative to growths in which substrate was withheld. Growth and expression conditions used are reported in **Supplementary Information Section V: Plate reader-based expression assay of sfGFP variants**.

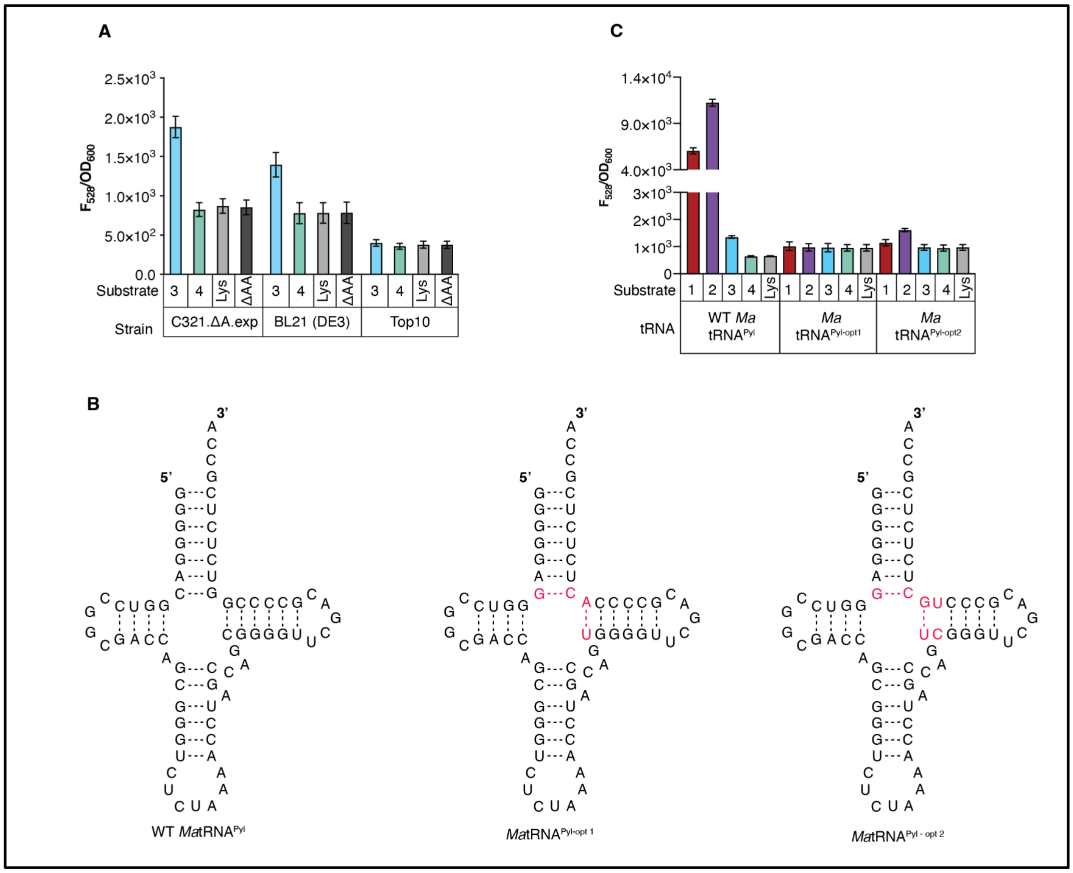

**Supplementary Figure 6. Effect of cell line and tRNA identity on the expression of sfGFP-3TAG in the presence of monomers 1-4.** (A) Shown is the $F_{528}/OD_{600}$ signal of C321.ΔA.exp, BL21 (DE3), and Top10 *E. coli* cells harboring pMega-*Ma*PylRS-*Ma*tRNA$^{Pyl}$ and pET22b-sfGFP-3TAG supplemented with 0.1 mM *(S)*-β$^{2}$-OH **3** or *(R)*-β$^{2}$-OH **4,** 24 hours after induction with 1 mM IPTG. Under these conditions the strongest $F_{528}/OD_{600}$ signal is observed in C321.ΔA.exp cells. (B) Sequences of tRNA$^{Pyl}$ variants evaluated in this work. *Ma*tRNA$^{Pyl\text{-}opt1}$ is a variant of *Ma*tRNA$^{Pyl}$ containing four mutations (red) present in *M. barkeri* tRNA$^{Pyl\text{-}opt}$.[14] *Ma*tRNA$^{Pyl\text{-}opt2}$ is a variant of *Ma*tRNA$^{Pyl}$ containing six mutations (red) present in *E. coli* tRNA$^{Sec}$.[15] (C) Shown is the $F_{528}/OD_{600}$ signal of C321.ΔA.exp *E. coli* harboring a pMega-*Ma*PylRS plasmid and pET22b-sfGFP-3TAG supplemented with 0.1 mM of monomers **1-4,** 24 hours after induction with 1 mM IPTG. The pMega-*Ma*PylRS plasmid carries either wild-type *Ma*tRNA$^{Pyl}$, *Ma*tRNA$^{Pyl\text{-}opt1}$, or *Ma*tRNA$^{Pyl\text{-}opt2}$.

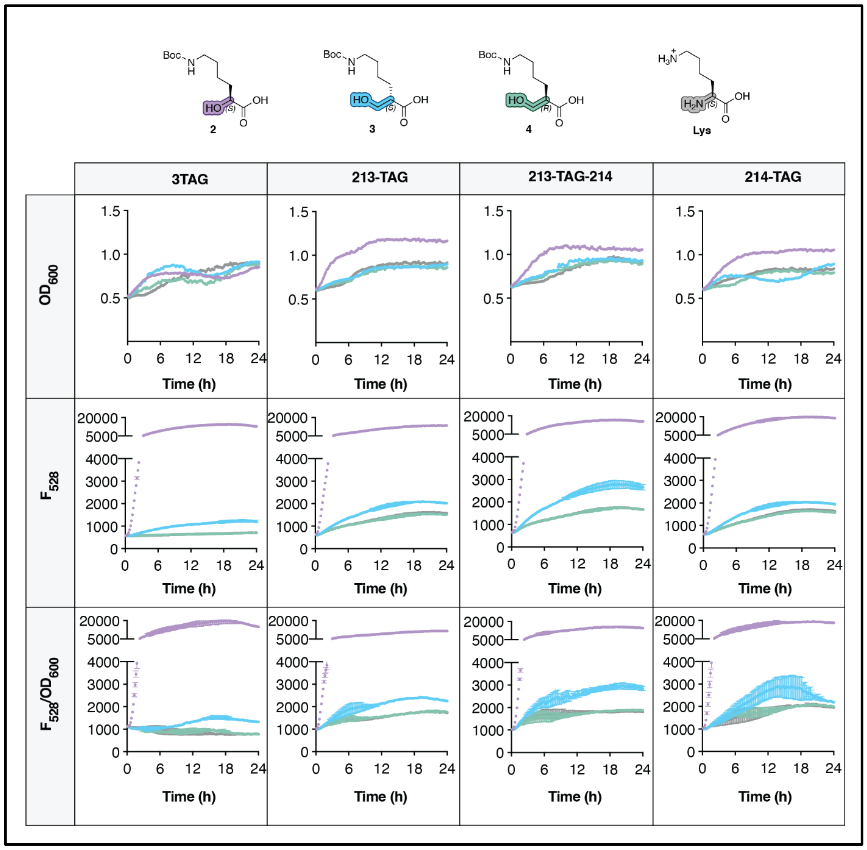

**Supplementary Figure 7. *Ma*PylRS supports incorporation of (*S*)-β[2]-OH-BocK into sfGFP at positions other than position 3.** Shown are growth ($OD_{600}$), GFP fluorescence ($F_{528}$), and growth-corrected ($F_{528}/OD_{600}$) curves for C321.ΔA.exp cells supplemented with 0.1 mM of monomer **2** - **4** or lysine (negative control), grown in a 96-well plate and monitored using an Agilent BioTek Synergy H1 Multi-Mode Microplate Reader. Increases in $F_{528}/OD_{600}$ signal in wells supplemented with monomer **3** relative to **4** or lysine suggest additional permissive sites for (S)-β[2]-OH-BocK incorporation into sfGFP. (n = 2, biological replicates).

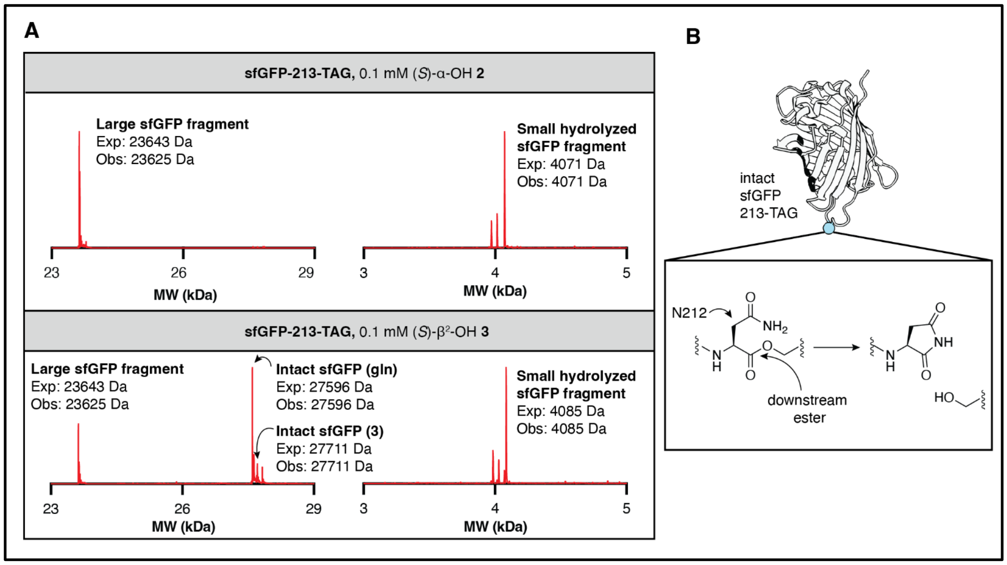

**Supplementary Figure 8. LC-HRMS characterization of sfGFP-213-TAG.** (A) Shown are deconvoluted mass spectra of purified sfGFP isolated from C321.ΔA.exp, *E. coli* transformed with pMega-PylRS and sfGFP-213-TAG and supplemented with either 0.1 mM (*S*)-α-OH **2** or (*S*)-$\beta^2$-OH **3**. Protein expressed with monomer **2** is obtained exclusively as the spontaneous hydrolysis product (large fragment: 23625.61 Da, small fragment: 4071.86 Da). Protein expressed in the presence of monomer **3** is obtained primarily as an analogous hydrolysis product (large: 23635.43 Da, small: 4085.75 Da). A small amount of protein material with a molecular weight corresponding to that of intact β-ester containing sfGFP-213-TAG is obtained (27711.73 Da). sfGFP containing glutamine instead of monomer **3** is the major contaminant (27596.14 Da). (B) The observed large hydrolysis fragment for sfGFP-213-TAG bearing either an α or β ester is 18 Da smaller than predicted, likely due to autohydrolysis by the asparagine upstream of the ester, formation of the succinimide and loss of water in the large fragment.[16]

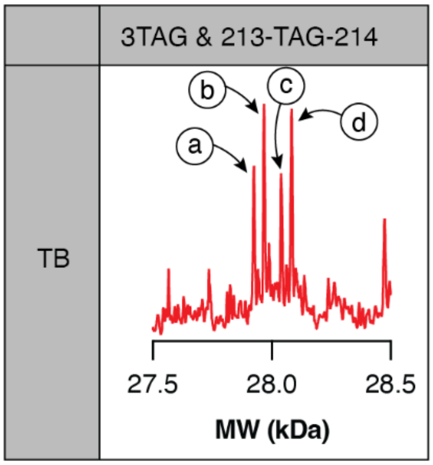

| Media | Prot. | Pos. 3 | Pos. 213-TAG -214 | MW (exp) | MW (obs) |
|---|---|---|---|---|---|
| TB | a | gln | gln | 27924 Da | 27925 Da |
| | b | **3** | gln | 27968 Da | 27968 Da |
| | c | gln | **3** | 28039 Da | 28041 Da |
| | d | **3** | **3** | 28083 Da | 28084 Da |

**Supplementary Figure 9. LC-MS characterization of sfGFP-3TAG-213-TAG-214 from C321.ΔA.exp, *E. coli* grown in terrific broth (TB).** Shown is the deconvoluted mass spectrum of purified sfGFP isolated from C321.ΔA.exp, *E. coli* transformed with pMega-PylRS and pET22b-sfGFP-3-TAG-213-TAG-214 and supplemented with 0.1 M *(S)*-$\beta^2$-OH **3**. The four major peaks in the spectrum correspond to sfGFP isoforms containing either: (a) Gln at position 3 and between positions 213 and 214; (b) *(S)*-$\beta^2$-OH **3** at position 3 (with residues 1 and 2 lost *via* hydrolysis) and Gln between positions 213 and 214, (c) Gln at position 3 and *(S)*-$\beta^2$-OH **3** between positions 213 and 214; and (d) *(S)*-$\beta^2$-OH **3** at both position 3 (with residues 1 and 2 lost *via* hydrolysis) and between positions 213 and 214.

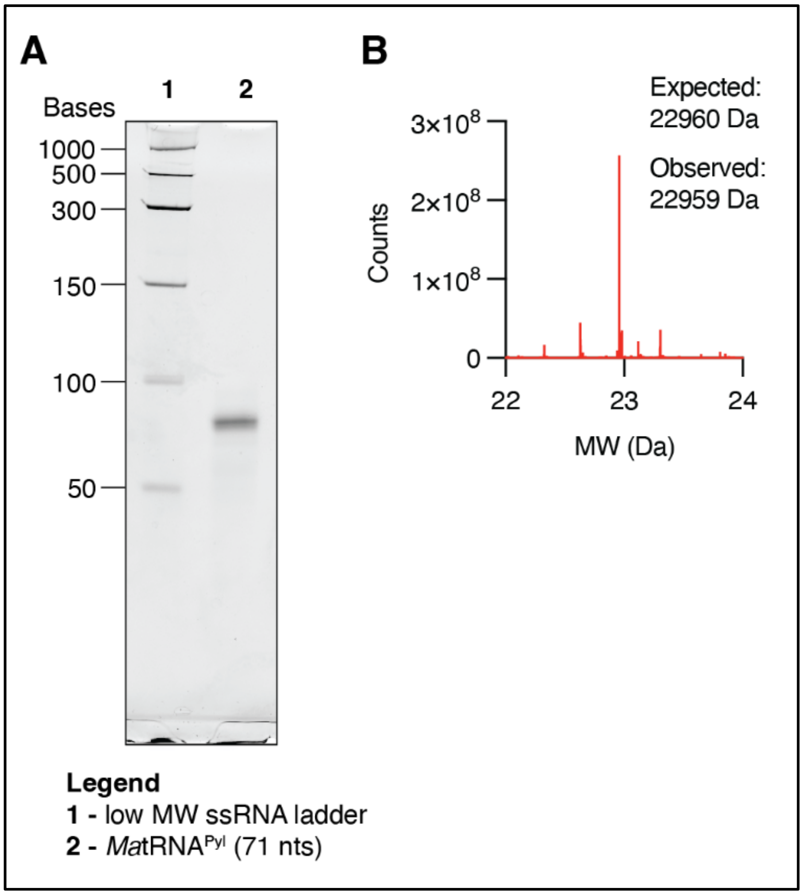

**Supplementary Figure 10. Characterization of *Ma*tRNA$^{Pyl}$ produced by transcription *in vitro*.** (A) 10% Mini-PROTEAN® TBE-Urea Gel loaded with a low range ssRNA ladder (NEB) (lane 1) and 100 ng *Ma*tRNA$^{Pyl}$ prepared using T7 RNA polymerase *in vitro* (see Supplementary Information for experimental details). The gel was subject to electrophoresis at 120 V for 1.5 h. (B) Deconvoluted mass spectrum of *Ma*tRNA$^{Pyl}$. The major peak corresponds to the expected MW of full length *Ma*tRNA$^{Pyl}$ with a single 5'-phosphate group. Lower MW bands correspond to early termination (n-1 and n-2) products.

VIII. Supplementary Tables 1 - 2

##### A. Table 1: DNA oligos used

| | Oligo Name | Sequence | Notes |
|---|---|---|---|
| 1 | MaPylT - F | CTAATACGACTCACTATAGGGGGACGGTCCGGCGACCAGCGGGTCTCTAAAACCTAGCCA | |
| 2 | MaPylT - R | TmGGCGAGAGACCGGGGCGTCGAACCCCGCTGGCTAGGTTTTAGAGACCCGCTGGTCGCCG | |
| 3 | MaPyl opt1 - F | cctagccagtggggttcgacgccccacTCTCTCGCCAAATTCGAAAAG | |
| 4 | MaPyl opt1 - R | ttttagagacccgctggtcgccggacccTCCCCCGTTCAGATGGTT | |
| 5 | MaPyl opt2 - F | cctagccagtcgggttcgacgccctgcTCTCTCGCCAAATTCGAAAAG | |
| 6 | MaPyl opt2 - R | cctagccagtggggttcgacgccccacTCTCTCGCCAAATTCGAAAAG | Same seq as MaPyl opt1 - R |
| 7 | sfGFP-E213TAG - F | AGATCCCAACtagAAGCGTGACC | |
| 8 | sfGFP-E213TAG - R | TTCGAAAGGACAGATTGTG | |
| 9 | sfGFP-E213TAGK214 - F | tagAAGCGTGACCACATGGTC | |
| 10 | sfGFP-E213TAGK214 - R | TTCGTTGGGATCTTTCGAAAG | |
| 11 | sfGFP-K214TAG - F | TCCCAACGAAtagCGTGACCACA | |
| 12 | sfGFP-K214TAG- R | TCTTTCGAAAGGACAGATTGTGTC | |

#### B. Table 2: Recipe for defined ΔgIn media

| Reagent | Final Concentration | Volume for 100 mL of ΔgIn media |
|---|---|---|
| 10% glycerol | 1% (v/v) | 10 mL |
| 50x M salts* | 1x | 2 mL |
| $MgSO_4$ | 5 mM | 200 µL |
| 5% Asp** | 0.25% (w/v) | 5 mL |
| 4mg/mL Leu | 80 µg/mL | 2 mL |
| 5x AA mix*** | 1x (400 µg/mL) | 20 mL |
| Trace metals* | - | 20 µL |
| MilliQ $H_2O$ | - | Up to 100 mL |
| *Described in Ref. 9 | | |
| **pH adjusted to 7.5 with NaOH | | |
| ***2 mg/mL of each amino acid except for Tyr, Cys, and Gln | | |
